# Predicting purification process fit of monoclonal antibodies using machine learning

**DOI:** 10.1101/2024.08.05.606711

**Authors:** Andrew Maier, Minjeong Cha, Sean Burgess, Amy Wang, Carlos Cuellar, Soo Kim, Neeraja Sundar Rajan, Josephine Neyyan, Rituparna Sengupta, Kelly O’Connor, Nicole Ott, Ambrose Williams

## Abstract

In early stage development of therapeutic monoclonal antibodies, assessment of ease of purification process development typically requires extensive experimentation. However, upstream protein expression and downstream purification experiments are often in conflict with timeline pressures and material constraints, limiting the number of molecules and process conditions that can reasonably be assessed. Recently, high-throughput batch-binding screen data along with improved molecular descriptors have enabled development of robust quantitative structure-property relationship (QSPR) models which predict monoclonal antibody chromatographic binding behavior from the amino acid sequence. This work describes a QSPR strategy for *in silico* monoclonal antibody purification process fit assessment. Principal Component Analysis is applied to extract a one-dimensional basis for comparison of molecular chromatographic binding behavior from multi-dimensional high-throughput batch-binding screen data. Kernel Ridge Regression is employed to predict the first principal component for new molecular sequences. This workflow is demonstrated with a set of 97 monoclonal antibodies for five chromatography resins in two salt types across a range of pH and salt concentrations. Model development benchmarks four descriptor sets from biophysical structural models and protein language models. The investigation illustrates the value QSPR models can provide to purification process fit assessment, and selection of resins and operating conditions from sequence alone.

## 0 Introduction

### 0.1 Experimental purification process fit assessment

As monoclonal antibodies (mAbs) have become a mature therapeutic modality, biopharmaceutical companies increasingly strive to streamline and accelerate manufacturing process development in order to bring new treatments to patients more quickly and at lower cost. In the mAb development pipeline, following discovery and humanization of molecular candidates, evaluation of the ease of development of therapeutic candidates (a.k.a. developability assessment) is crucial in ensuring rapid and smooth development timelines. Developability assessment aims to identify candidates with molecular properties that minimize a portfolio of risks including pharmacokinetic clearance, polyspecificity, physicochemical stability (oxidation, deamidation, isomerization), physical stability (solubility, viscosity, aggregation propensity), and, most relevant to the present work, manufacturability^1^. Traditionally, developability assessment involves a set of empirical assays; however, the material requirements for direct study often delays the timeliness of assessment and limits the number of molecular candidates and process conditions which can realistically be assessed.

The evaluation of mAb manufacturability for downstream purification, or purification process fit assessment, is primarily focused on evaluation of chromatographic binding behavior. In large-scale purification of mAbs, use of platform purification processes has become a well-established practice^2,3^. Use of platform purification processes for molecules with similar properties leverages platform knowledge to reduce downstream bioprocess development work, mitigate development risks, ensure process fit in manufacturing facilities, and accelerate development timelines. Assessment of chromatographic binding behavior aims to identify atypical molecules which might not readily fit within platform purification processes, evaluate available binding mechanisms and process levers, guide resin selection, and ascertain robust separation process conditions. Historically, a common strategy to assess chromatographic binding behavior has been measurement of molecular retention time during gradient elution^4^. However, gradient retention time experiments are not easily parallelizable, typically only evaluate binding at a single pH, and are not readily applicable to all modes of chromatography.

In order to overcome the limitations of gradient retention time experiments, purification process fit assessment commonly relies on high-throughput batch-binding screens^5–9^. These screens measure equilibrium binding of a molecule to a set of chromatography resins across a wide range of pH and salt concentrations. Automated workflows debottleneck data generation by screening multiple binding conditions in parallel in 96 or 384-well plates. The set of resins screened typically include ligands with diverse combinations of charged and hydrophobic moieties. While high-throughput batch-binding screens have greatly accelerated assessment of chromatographic binding behavior, they are still limited in throughput and require considerable time investment in upstream protein expression at a point early in process development when cell culture titers may be low and material scarce. *In silico* quantitative structure-property relationship (QSPR) models for prediction of chromatographic binding behavior are needed in order to further accelerate and debottleneck purification process fit assessment and inform and de-risk purification process development decisions.

### 0.2 *In silico* purification process fit assessment

While there has been considerable research and industrial application of biophysical structural descriptors reflecting many of the mechanisms underlying mAb developability risks^10–14^, the application of *in silico* models to prediction of mAb chromatographic binding behavior is less ubiquitous. QSPR models leverage molecular descriptors to predict experimental results.

Early QSPR models of preparative-scale chromatographic binding behavior utilized small non-mAb proteins. Initial studies focused on prediction of small protein chromatographic retention (the amount of salt required for gradient elution) on multiple modes of chromatography including cation exchange chromatography^15–17^ (CEX), anion exchange chromatography^18^ (AEX), hydrophobic interaction chromatography^19–21^ (HIC), and multimodal chromatography including multimodal anion exchange chromatography^22^ (MMAEX) and multimodal cation exchange chromatography^17^ (MMCEX). For CEX of small proteins, prediction of mechanistic model adsorption isotherm parameters^23,24^ has also been studied^25,26^. Predicted isotherm parameters can subsequently be used for prediction of high-throughput batch-binding screens, gradient retention times, or other chromatographic behavior. Prior QSPR studies for small proteins generally share similar modeling strategies: biophysical descriptors based on crystal structures (e.g., pH-dependent electrostatics, lipophilicity, surface hydrophobicity, etc.), a small training set (<30 molecules) and small randomly withheld test set (∼2 molecules), and k-fold cross validation (CV).

Development of QSPR models of preparative scale chromatographic binding behavior for larger multi-chain molecules such as mAbs has been more limited. Robinson et al^27^ predicted chromatographic retention across multiple chromatographic modalities (CEX, MMCEX, and HIC) with a training set of 20 fragment antigen-binding regions (Fabs). They leveraged global Molecular Operating Environment^28^ (MOE) homology_-_model derived structural descriptors, surface patch descriptors based on clusters of solvent accessible aliphatic or aromatic residues, and local descriptors capturing changes in local spatial aggregation propensity^29^ (SAP) and electrostatic potential due to point mutations. At the Highland Games^30^, an academic and industrial QSPR model benchmarking competition, the prediction of chromatography retention time on ion exchange chromatography (Capto Q, POROS 50HS) category was won using a training set of 28 mAbs, homology model-based structural descriptors, k-fold cross validation, and a partial least squares (PLS) regressor. Second-place had success predicting retention time with a Steric Mass Action (SMA) model^31^, assuming a characteristic charge calculated from amino acid pKa values. Recently Saleh et al^32^ reported prediction of CEX isotherm parameters with a training set of 21 large molecules using homology-model-based structures and structural descriptors from BioLuminate^10,33^. Diversity of the training set was further supplemented by digesting the IgG1 and IgG4 mAbs with papain, and by making independent predictions for each molecule at each pH level. The modeling strategy leveraged a two-step feature selection pipeline and a Gaussian Process Regressor with a nonlinear kernel^34^. Subsequently, Hess et al applied a similar strategy to prediction of gradient retention time^35^ and isotherm parameters^36^ on MMAEX chromatography with a dataset of 64 and 59 large molecules, respectively.

There is a significant opportunity to improve molecular descriptors and QSPR models for HIC. For their aforementioned QSPR model predictions of Fab Capto Phenyl retention time, Robinson et al^27^ noted a systematic over-prediction of HIC retention times and low predictive ability with a test set of 4 Fabs. While developing structural biophysical descriptors for prediction of HIC retention time on butyl-NP5 resin with the dataset of 137 mAbs tested by Jain et al^4^, Park et al^14^ report poor correlation even for the best descriptor identified. In their comparison of the correlation of structural biophysical descriptors to HIC retention time using four benchmark datasets, Waibl et al^11^ similarly reported poor results^37^. Both Park and Waibl found that choice of homology modeling software package and hydrophobicity scale have a major impact on the accuracy of hydrophobic descriptors, but saw no improvement from conformational sampling by Gaussian Accelerated Molecular Dynamics^38^ (GaMD). They reported the best correlations with hydrophobicity scales from Black-Mould^39^, Jain^4^, and Wimley-White^40^.

Despite the significant progress in QSPR modeling, existing methods may not fully meet the requirements of purification process fit assessment. While prediction of gradient retention time can be used to identify atypical molecules for a specific resin and pH condition, this method may not be as useful as high-throughput batch-binding screens for simultaneously informing selection of resin, mode of operation, and buffer pH and salt concentrations. While isotherm parameter prediction has been successfully demonstrated for CEX and MMAEX chromatography of mAbs, the methodology also has drawbacks. Isotherms typically have multiple parameters, thus molecular comparison must be performed in a high-dimensional basis which can limit interpretability. Furthermore, the accuracy of isotherm parameter values is dependent on the accuracy of the underlying assumptions of the selected isotherm. Batch-binding experiments are complex systems that may exhibit deviations from a theoretical model, potentially limiting the relevance of isotherm parameter predictions. During isotherm parameter estimation, “sloppy” parameter sensitivities ^41^ may result in large parameter uncertainties, or a large number of possible equally well-fitting parameter sets, undesirable properties for a QSPR prediction target. Alternatively, compression of high-throughput batch-binding screen data with Principal Component Analysis^42^ (PCA) allows for a low-dimensional representation of the actual measured binding behavior and is not limited by biases resulting from assumption of a specific structure for the adsorption dynamics.

### 0.3 Emergence of sequence-based protein language model descriptors

Recently, protein language models have emerged as a data-driven molecular representation strategy that may supplement the structural biophysical descriptors. Protein language models are trained in an unsupervised manner on large unlabeled publicly available protein sequence datasets^43^ to embed a molecular descriptor vector. Early protein language models^44–46^ used neural networks to embed short sequences (<250 amino acids). In 2017, invention of the attention mechanism in transformer models^47^ enabled application of protein language models to proteins with longer sequences (typically <1024 amino acids). Subsequently, there has been a proliferation of transformer-based protein language models trained on large protein sequence datasets ^48–50^.

Remarkably, protein language models trained in an unsupervised manner learn relevant structural properties of proteins. Transformers have been shown to learn inter-amino acid attention maps that reproduce protein contact maps comparably to state-of-the-art contact map prediction methods^51,52^. In light of this finding, several techniques have emerged that leverage protein language model embeddings for tasks including supervised prediction of protein secondary structure^53^, tertiary structure^50,54^, quaternary structure of mAb complimentary-determining regions ^55–57^ (CDRs), and cryogenic electron microscopy (cryo-EM) structure determination^58^. Given the broad utility of protein language model embeddings for modeling protein structure, it is probable they may be complementary to existing structural biophysical descriptors and beneficial to prediction of protein chromatographic binding behavior.

### 0.4 Improving upon prior methodology

This work presents a QSPR modeling strategy for assessment of mAb purification process fit from amino acid sequence. To simplify comparison of mAb chromatographic binding behavior, PCA is applied to compress high-throughput batch-binding screen data into a one-dimensional metric: the first principal component (PC1). QSPR models are trained to predict PC1 of high-throughput batch-binding screen data. Applicability of this modeling strategy is demonstrated with seven different combinations of chromatography resin and salt type. Models are trained with 87 mAbs, and tested with 10 mAbs selected at random with stratification by PC1 quantile. The work compares the accuracy of models trained using different types of descriptors: biophysical structural descriptors from commercial software packages, sequence-embeddings from open-source protein language models, and the combination of all of these descriptors. Given the large number of descriptors and relatively small number of molecules in the training set, the risk of overfitting is a key challenge. To address the risk of overfitting, the QSPR regression model leverages correlation-based feature filtering, model-based recursive elimination, and a regularization penalty during Kernel Ridge Regression^59^ (KRR). Our results demonstrate the utility of QSPR models for *in silico* mAb purification process fit assessment from amino acid sequence.

## 1 Methods

### 1.1 Overall Approach

QSPR models aim to predict the PC1 of batch-binding screen data from molecular descriptors. A graphical outline of model training and application is found in Figure 1.

**Figure 1.**
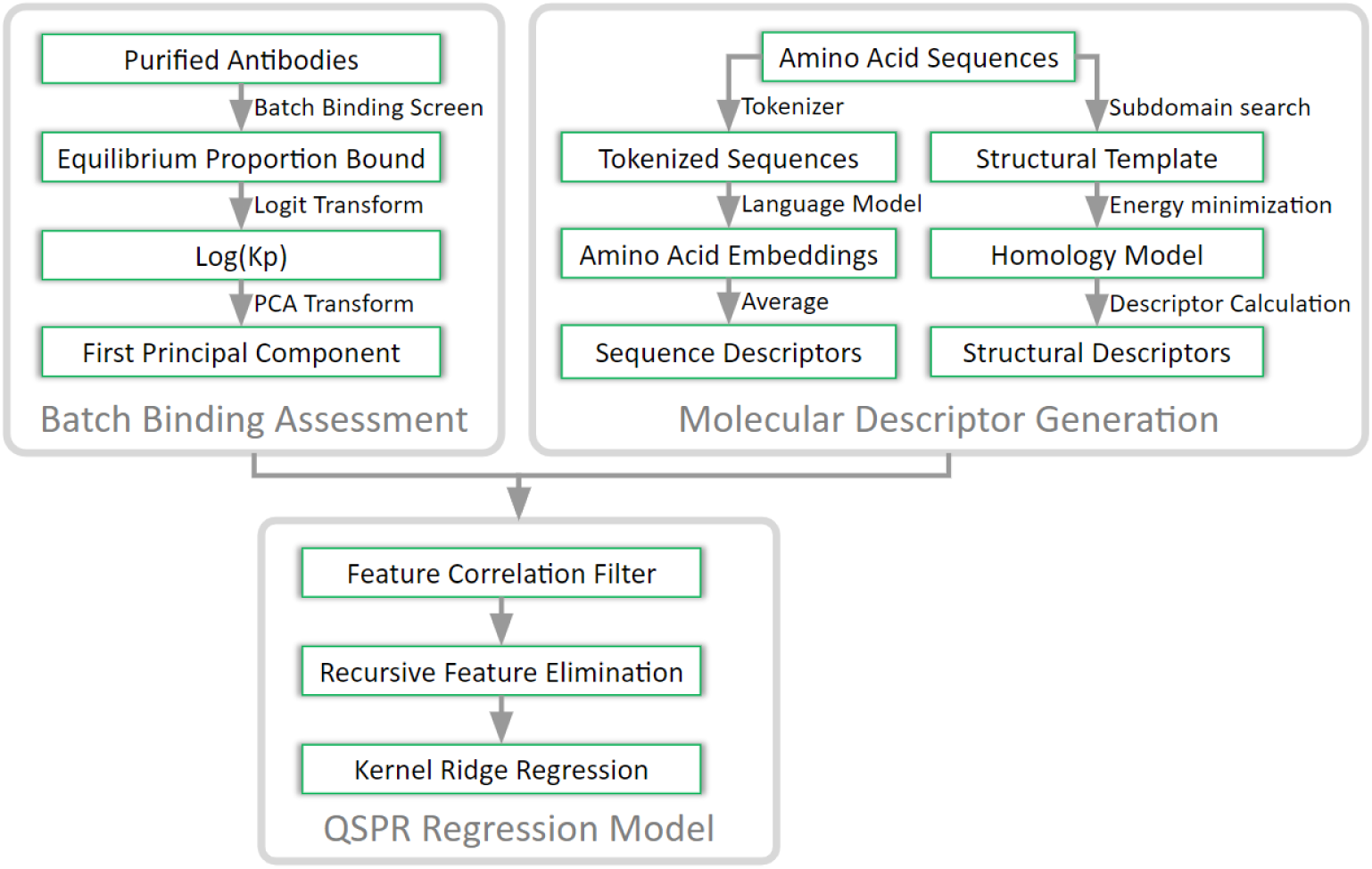
Graphical outline of QSPR model development.

### 1.2 High-Throughput Batch-Binding Screen

The dataset for model development and evaluation is composed of a set of 97 mAbs (90 IgG1, 2 IgG2, 5 IgG4). The mAbs are expressed in Chinese Hamster Ovary cells (Genentech, Inc.). Purification is performed in a two-step process. First, mAbs are captured by affinity chromatography with Protein A resin. Next, preparative-scale size exclusion chromatography is used to separate the mAb monomer from aggregates and other size variants to >95% monomer purity. Purified mAbs are buffer exchanged and concentrated to ≥10 g/L (20 mM Tris acetate, pH 5.5) using 10 kDa Amicon Ultra-15 filters (EMD Millipore). Concentrated mAb feedstocks are diluted in order to reach the desired pH and salt concentration for each equilibrium binding condition prior to the high-throughput batch-binding screen.

High-throughput batch-binding screens are performed for five resins in two background salt species across the range of pH and salt conditions tested (24 combinations of pH and salt concentration) according to the methods described by McDonald^9^. The set of resins in the high-throughput batch-binding screens evaluate binding with different combination of charged and hydrophobic functional groups. Ion exchange resins (CEX and AEX) are tested in sodium acetate (NaOAc). The HIC resin is tested in sodium sulfate (Na_2_SO_4_), which is a kosmotrope and promotor of hydrophobic interactions^60^. Mixed mode resins are tested in both salt species to potentially observe different modes of protein binding. Details of the chromatography resins assessed and range of pH and salt conditions tested for each resin can be found in Table 1.

**Table 1.**
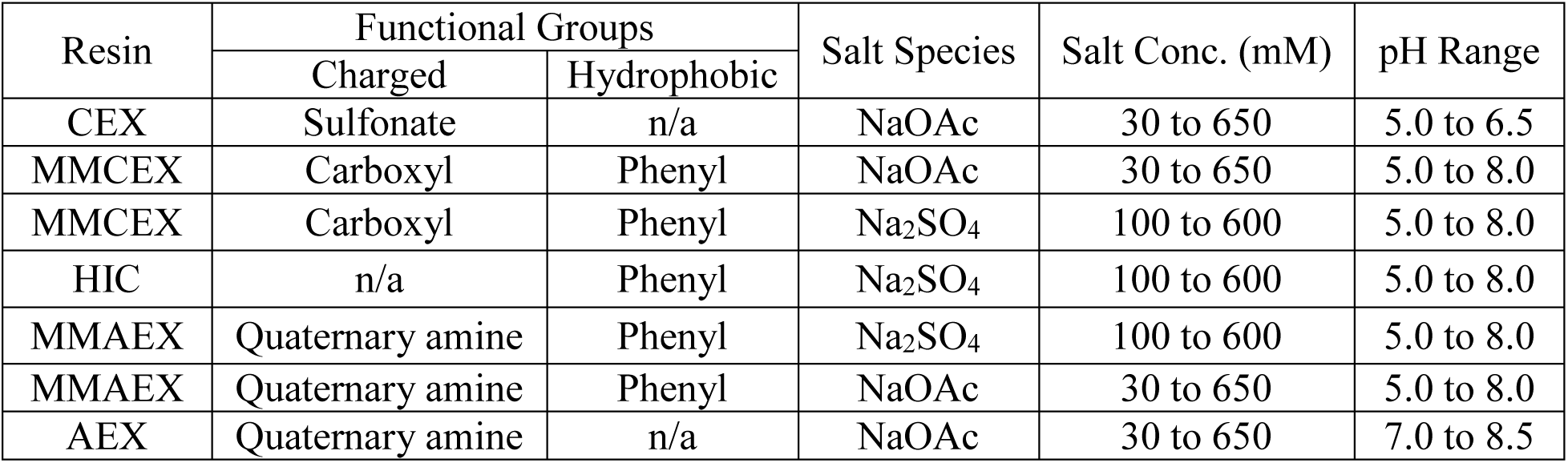
Molecular Batch-Binding Screen pH, salt, and resin conditions tested.

| Resin | Functional Groups |  | Salt Species | Salt Conc. (mM) | pH Range |
| --- | --- | --- | --- | --- | --- |
|  | Charged | Hydrophobic |  |  |  |
| CEX | Sulfonate | n/a | NaOAc | 30 to 650 | 5.0 to 6.5 |
| MMCEX | Carboxyl | Phenyl | NaOAc | 30 to 650 | 5.0 to 8.0 |
| MMCEX | Carboxyl | Phenyl | Na<sub>2</sub>SO<sub>4</sub> | 100 to 600 | 5.0 to 8.0 |
| HIC | n/a | Phenyl | Na<sub>2</sub>SO<sub>4</sub> | 100 to 600 | 5.0 to 8.0 |
| MMAEX | Quaternary amine | Phenyl | Na<sub>2</sub>SO<sub>4</sub> | 100 to 600 | 5.0 to 8.0 |
| MMAEX | Quaternary amine | Phenyl | NaOAc | 30 to 650 | 5.0 to 8.0 |
| AEX | Quaternary amine | n/a | NaOAc | 30 to 650 | 7.0 to 8.5 |

The output of the high-throughput batch-binding screens is equilibrium proportion of protein bound for each pH, salt, and resin condition tested. Proportion bound data is sometimes also reported as a partition coefficient (Kp, Equation 1). Prior to modeling, a logit transform is applied to batch binding proportion bound data^9^.

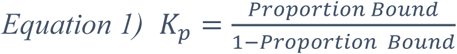

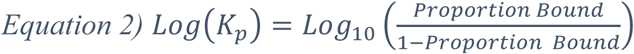

The logit transform linearizes the proportion bound data, which follows a sigmoidal form, from an output range of [0,1] to [-∞, +∞]. Logit transformed values are then truncated to the range [-0.5, 2.0], approximately [0.24, 0.99] as a proportion bound, to eliminate the impact of noisy measurement of proportion bound at the extremes. Use of the logit transform enables the application of linear models to predict the Log(Kp) values. Contour plots of Log(Kp) values across pH and salt conditions are generated in Plotly^61^ following cubic spline interpolation with SciPy^62^. In order to simplify the comparison of chromatographic binding behavior across molecules, PCA is applied to the set of standardized Log(Kp) values (mean centered, scaled to unit variance). For a given resin and salt type, PCA is applied across pH levels and salt concentrations. The output is a one-dimensional metric (PC1) representing binding of each molecule for a given resin and salt type. The PC1 of this data structure captures the direction of maximal covariance within the set of binding conditions tested. Standardization and PCA are implemented using Scikit-Learn^59^. For visualization purposes, PC1 values have been oriented so that a larger value corresponds to stronger binding: for cases where the Pearson correlation between PC1 and average high-throughput batch-binding screen Log(Kp) value for the set of molecules is negative, the PC1 values are multiplied by negative one. This manipulation does not impact the meaning or interpretation of PC1 values.

### 1.3 Molecular Descriptor Generation

Full-mAb homology models are generated for each molecule using the automatic Antibody Modeler in the Molecular Operating Environment^28^ software package^28^ using the Immunoglobulin (Ig) Model Type. Antibody Modeler searches the Protein Data Bank database for subdomain structural templates by length and sequence similarity. The best homology model is selected by structure score, which is a function of backbone topology, Ramachandran phi/psi probability, crystallographic occupancies, and temperature factors as described in by Maier et al^63^. The final model structure is obtained via energy minimization of the highest scoring homology model leveraging the EHT force field in Amber 10^28^. The framework region templates used in the homology model of each molecule^64^ are attached in Supplemental Table S1. Comparison of QSPR dataset sequences and corresponding framework region templates found no molecules to be sequence-identical in both Fab heavy and light chain.

The homology model-based biophysical descriptors are generated using two *in silico* software tools: the Enhanced Protein Properties and Patches tool in BioMOE^65^ and the Calc_Protein_Descriptors module in Maestro BioLuminate^10,33,66^. Descriptors are generated at pH 5, 6, 7, and 8. To reduce feature multicollinearity, descriptors at pH 5, 7, and 8 are only retained if they exhibit non-zero variance across the pH levels for all molecules in the training dataset. BioMOE is used to generate 248 descriptors (72 descriptors at pH 6 only and 44 descriptors at pH 5, 6, 7, and 8). BioLuminate is used to generate 1626 descriptors (878 descriptors at pH 6 only and 187 descriptors at pH 5, 6, 7, and 8). A summary of the pH levels of BioMOE and BioLuminate descriptors generated may be found in Supplemental Table S2 and S3 respectively. Model development utilized full-mAb structural descriptors based on preliminary analysis finding PC1 values have higher average correlation to full-mAb structural descriptors than Fv descriptors for the 30 most highly correlated BioLuminate descriptors (Supplemental Figure S1).

Two sets of sequence-based descriptors (sequence embeddings) are generated using pre-trained protein language models. ProtTrans^49^ (ProtT5-XL-UniRef50) is used to generate a 1024-dimension sequence embedding. ESM-2^50,67^ (esm2_t6_8M_UR50D) is used to generate a 320-dimension sequence embedding. While the ProtTrans embedding dimension is fixed, various sizes of model with different embedding dimensions are available for ESM-2. Preliminary analysis does not find larger dimension ESM-2 sequence embeddings to have significantly higher correlation to PC1 values (Supplemental Figure S1). To reduce the risk of overfitting in a high-dimensional descriptor space, the ESM-2 version with the smallest embedding dimension is used. A matrix of amino acid embedding vectors (size *m* embedding size by *n* sequence amino acids) is generated for the heavy and light chain of each molecule. For each molecule, heavy and light chain amino acid embedding vectors are concatenated along the sequence dimension. Molecular descriptors (an average molecular embedding vector) are generated for each molecule by averaging across the amino acid dimension for the concatenated amino acid embedding vectors for each molecule, resulting in a size m descriptor vector for each molecule.

### 1.4 Machine Learning

An overview of the machine learning pipeline including data splits, feature selection, and model optimization is shown in Figure 2.

**Figure 2.**
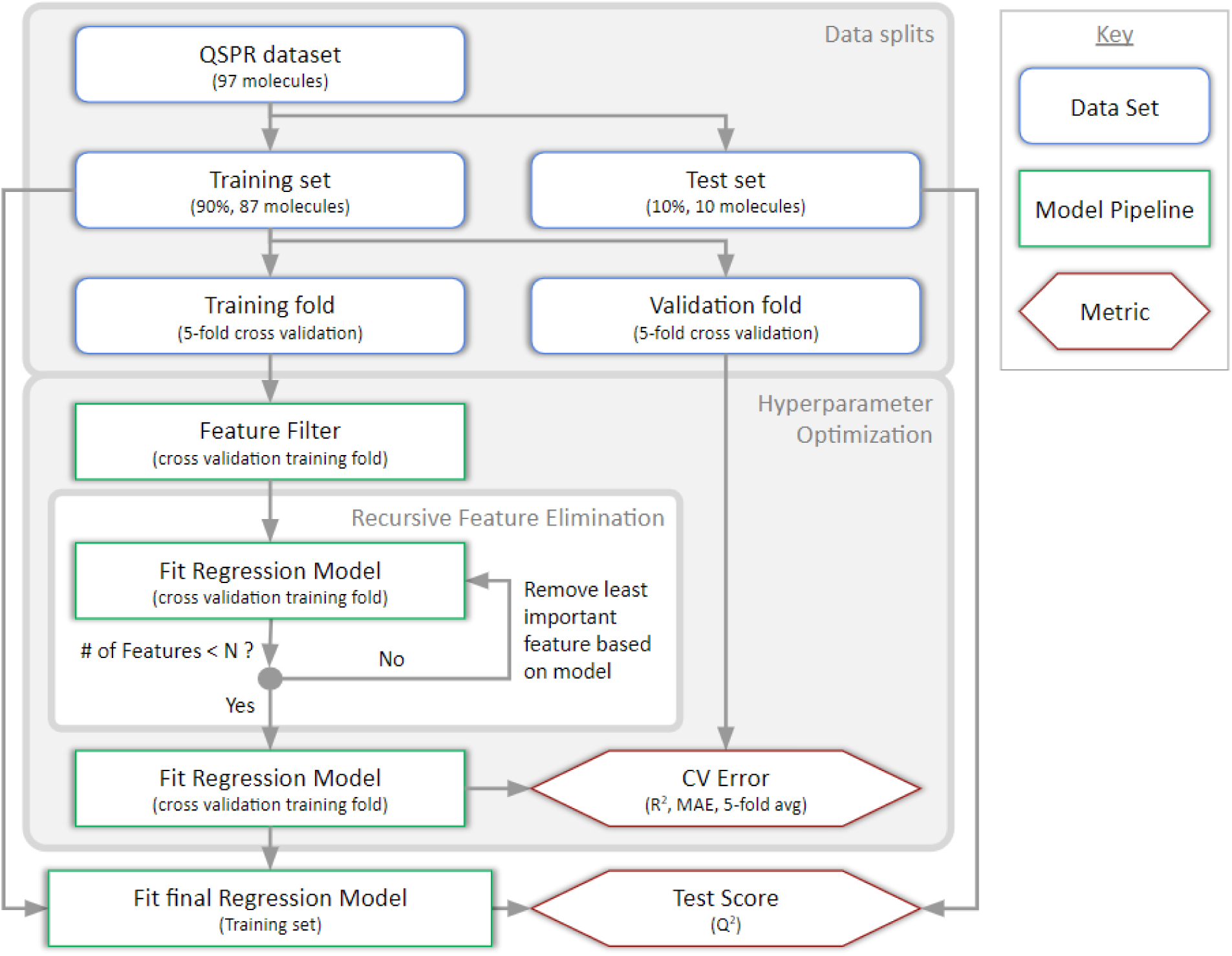
Summary of regression pipeline and cross-validation strategy. Hyperparameter optimization is performed iteratively via Bayesian Optimization. The best hyperparameter set is used when fitting the final regression model.

A total of 97 molecules are used for model training and evaluation. For each resin/salt combination, molecules are binned into seven quantiles based on their PC1 value. Train/test splits and 5-fold cross validation splits are performed randomly with stratification based on PC1 quantile to ensure diverse and rigorous test/validation sets. This procedure results in the test set containing different molecules for each resin and salt type modeled. For each model, the withheld test set consists of 8-10 mAbs (8-10% of the dataset). Reduced test set sizes for some models were a result of a post-hoc analysis of sequence similarity between the training and test sets (Supplemental Figure S2). Molecules with high sequence similarity to the training set (sequence identity >95% ^13,68^) were removed from the test set to ensure robustness of the reported model accuracy. During cross validation the randomly selected training and validation set consist of 69 (80% of the training set) and 18 (20% of the training set) molecules, respectively. Data splits were performed with Scikit-Learn^59^.

Hyperparameter optimization of Scikit-Learn model pipelines is performed by Bayesian Optimization with default algorithm parameter values using Scikit-Optimize^69^. Model pipeline hyperparameter ranges for Bayesian Optimization are found in Supplemental Table S4. The Bayesian Optimization routine is initialized with 100 pseudo-random model hyperparameter sets selected by Latin Hypercube Sampling. Subsequently 100 iterations of Bayesian Optimization are performed, each evaluating the model pipeline with a single hyperparameter set. The number of hyperparameter sets was chosen as a tradeoff between model performance and computational time. Hyperparameter sets are scored by Mean Absolute Error (MAE) of the fit regression model on the cross validation set averaged across the cross validation folds. Models are scored by the coefficient of determination, which is referred to as R^2^ for the training set and Q^2^ for the test set for clarity.

Machine learning tasks with a higher number of predictor variables than the number of samples in the dataset face a risk of overfitting. To address the risk of overfitting in a high-dimensional descriptor space, a two-part feature selection pipeline is applied. Prior to training regression models, all features are standardized (by mean centering and unit variance scaling). The feature selection pipeline initially leverages the K-Best filter method, which selects features with the highest Pearson correlation coefficient. Features are subsequently further reduced using recursive feature elimination (RFE), which iteratively eliminates features with the lowest feature importance in the model. Following RFE, the final models include less than 30 features. The exact number of features selected by Pearson correlation and RFE is determined by Bayesian Optimization. Filtering features prior to model-based RFE removes non-predictive features, reduces the number of iterations (and computational time) required for RFE, and should improve the accuracy of model feature importance scores and consequently the robustness of RFE. RFE and the final regression model rely on slightly different regressors. RFE uses a linear Ridge Regression (RR) model. Features selected by RFE are then inputs to Kernel Ridge Regression (KRR) in the final model. RR and KRR both rely on regularization, an extra penalization coefficients to the model cost function, to discourage model overfitting. The RR and KRR regularization penalties are optimized by Bayesian Optimization. Machine learning pipelines in this work are implemented with Scikit-Learn^59^.

## 2 Results

### 2.1 Training Set Molecule Diversity

In order to ensure development of robust models and reduce risk of data drift, it is important to consider whether the training set molecules are representative of the larger set of possible mAbs. Sequences for clinical stage mabs with international nonproprietary names (INN) are available Wilkinson et al.^70^ and Fv sequences are collated in the Therapeutic Structural Antibody Database^71^ (Thera-SAbDab). As most mAb sequence variation lies in the fragment variable domain (Fv), Fv descriptor distributions are useful for comparison of mAb dataset diversity. Since binding in the modes of chromatography studied (Table 1) is driven by electrostatic and hydrophobic forces, the distribution of these two molecular properties is relevant. Figure 3 compares the distributions of mAb Fv charge and hydrophobicity between the present QSPR training dataset and clinical stage mAbs in Thera-SAbDab.

**Figure 3.**
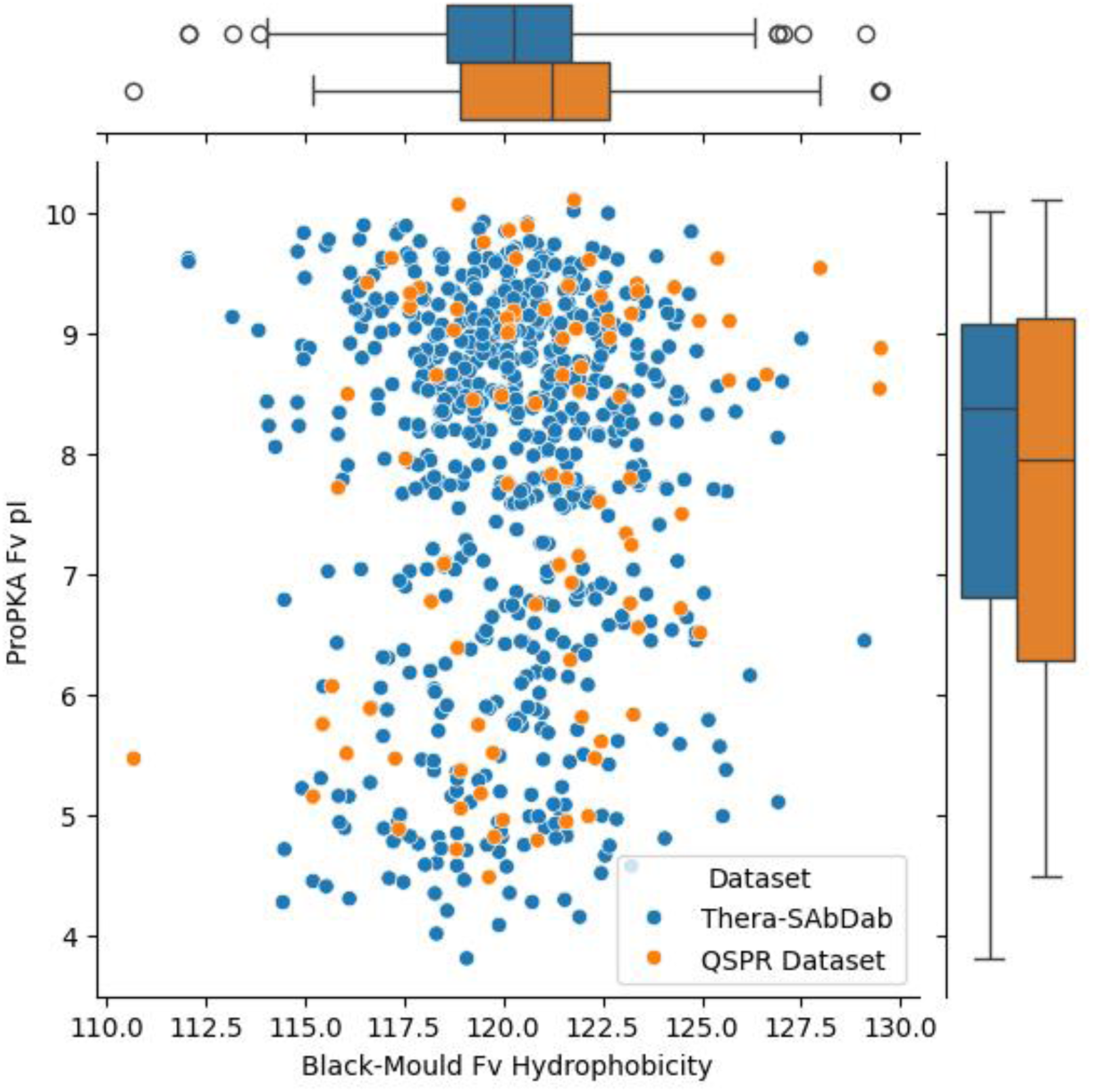
Comparison of the distribution of mAb Fv domain charge (ProPKA isoelectric point) and hydrophobicity (sum of residue Black-Mould hydrophobicity) for the present QSPR Dataset vs the Thera-SAbDab.

Charge and hydrophobicity respectively are represented by BioLuminate descriptors for non-ideal isoelectric point (pI) from ProPKA^72^ and sum of residue Black & Mould hydrophobicity^39^, which has been found to correlate well to mAb HIC retention time^11,14^. Each point represents a single molecule, while the comparability of molecular property distributions is illustrated by the boxplots along each axis. The similarity in the distribution of biophysical properties between the QSPR dataset used in this investigation and the Thera-SAbDab dataset suggest that the QSPR distribution is representative of clinical stage mAbs and should be relevant to prediction of chromatographic binding behavior for future molecules going through purification process fit assessment.

### 2.2 Dimensionality Reduction of Experimental Data with PCA

To simplify comparison of chromatographic binding behavior across molecules, PCA is applied to the high-throughput batch-binding screen data. PC1, as it captures the direction of maximal covariance in the dataset, serves as a representative one-dimensional metric of molecular chromatographic binding behavior. The proportion of variance explained by principal components is shown in Figure 4A. A visualization of reconstructed CEX Log(Kp) values after taking the inverse PCA transform for each PC1 value is shown in Figure 4B.

**Figure 4.**
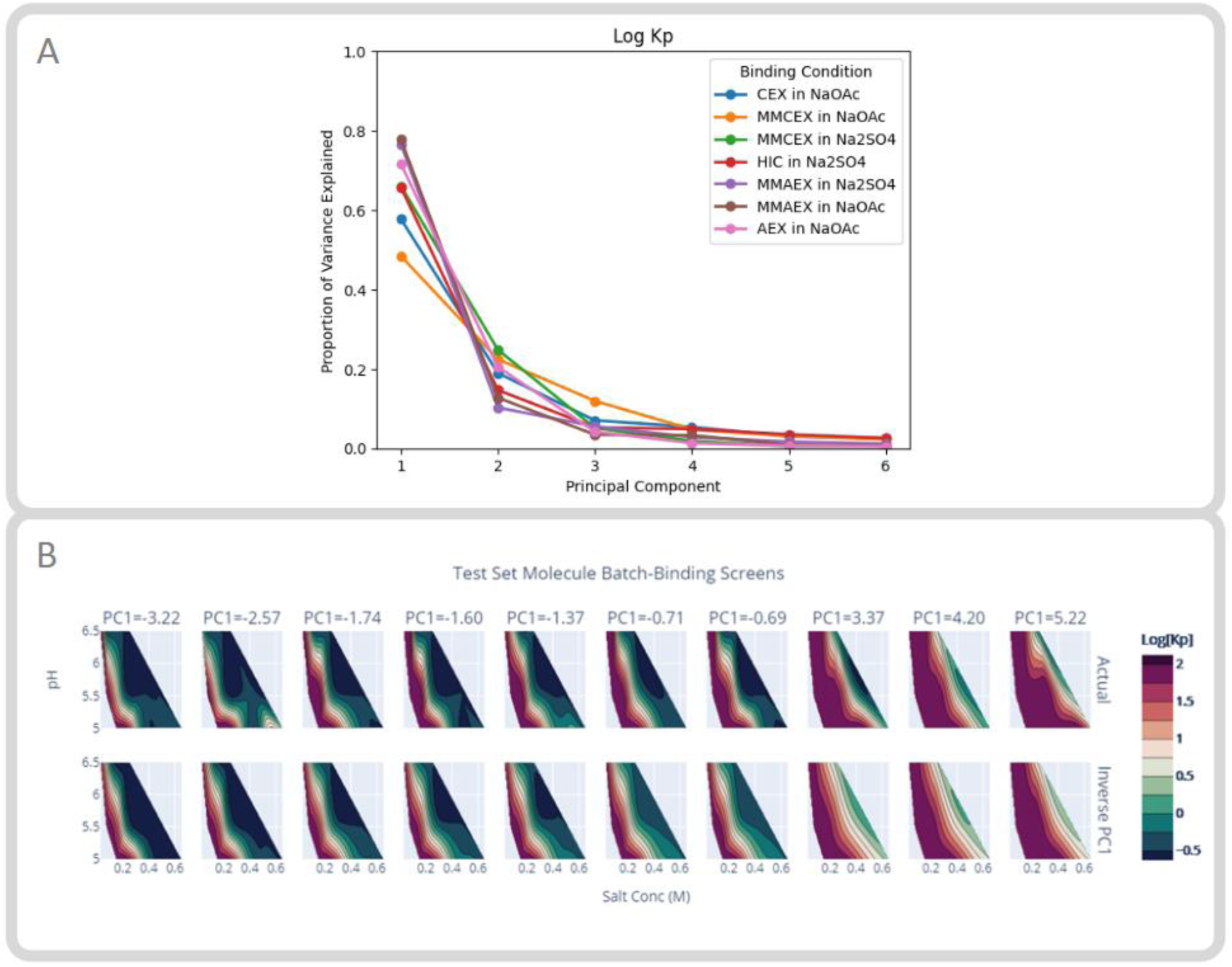
Application of PCA to high-throughput batch-binding screen data. (A) The Scree Plot illustrates the proportion of variance explained by PCA for molecular high-throughput batch-binding screen data. (B) Contour plots of actual Log(Kp) values vs Log(Kp) values from inverse PCA transform of PC1 values for CEX in NaOAc test set molecules. Figures similar to Figure 4B for every combination of resin and salt type studied may be found in Supplemental Figure S3.

As illustrated in Figure 4A, PC1 accounts for the majority of variance in batch binding data across all resin and salt conditions. The high proportion of variance explained by PC1 indicates a high degree of covariance among the data points across pH and salt concentrations tested for a given binding condition. For a quantitative visualization of how well PC1 represents Log(Kp) data, crossplots comparing actual Log(Kp) values to Log(Kp) values from the inverse PCA transform of PC1 values for every combination of resin and salt type studied may be found in Supplemental Figure S4. The inverse PCA transform is used to map predicted PC1 values back into the multi-dimensional experimental space in order to visualize Log(Kp) contour plots and inform purification process development decisions. In general, taking the inverse PCA transform of PC1 values results in Log(Kp) contour plots that are qualitatively similar to the actual Log(Kp) contour plots (Figure 4B, Supplemental Figure S4). Visual comparison in Figure 4B indicates that molecules with similar PC1 values have highly similar actual Log(Kp) contour plots (even for CEX in NaOAc, for which PC1 has a relatively low proportion variance explained). PC1 captures relative trends in chromatographic binding behavior across pH and salt concentrations for a given resin and salt type, therefore it is suitable for comparison of batch-binding screens across mAbs.

### 2.3 Model Accuracy

The QSPR model strategy leverages a final KRR model fit to features selected by recursive feature elimination with a RR model. RR and KRR both rely on regularization, an extra penalization coefficients to the model cost function, to discourage model overfitting. However, while RR is a linear regression technique KRR implements nonlinear regression by implicitly mapping data into a higher dimensional space through computationally efficient kernel functions. For data of our size, the expressivity of KRR improves model accuracy as compared to linear regression techniques. While prior works have mostly relied on kernel support vector regression (K-SVR) to learn non-linear functions, this work relies on KRR. The form of the model learned by K-SVR and KRR are identical, but KRR is faster to train on small datasets because it has a closed-form solution^59^.

Training with the described QSPR regression pipeline was successful for all binding conditions. For the best performing descriptor sets, model training accuracy (R^2^) for CEX, AEX, and MMAEX in NaOAc were all above 0.92, with the best performing descriptor sets for all other binding conditions reporting R^2^ between 0.78-0.84 (Supplemental Figure S5). A comparison of model performance on the withheld test set for each binding condition evaluated with each descriptor set is found in Table 2.

**Table 2.**
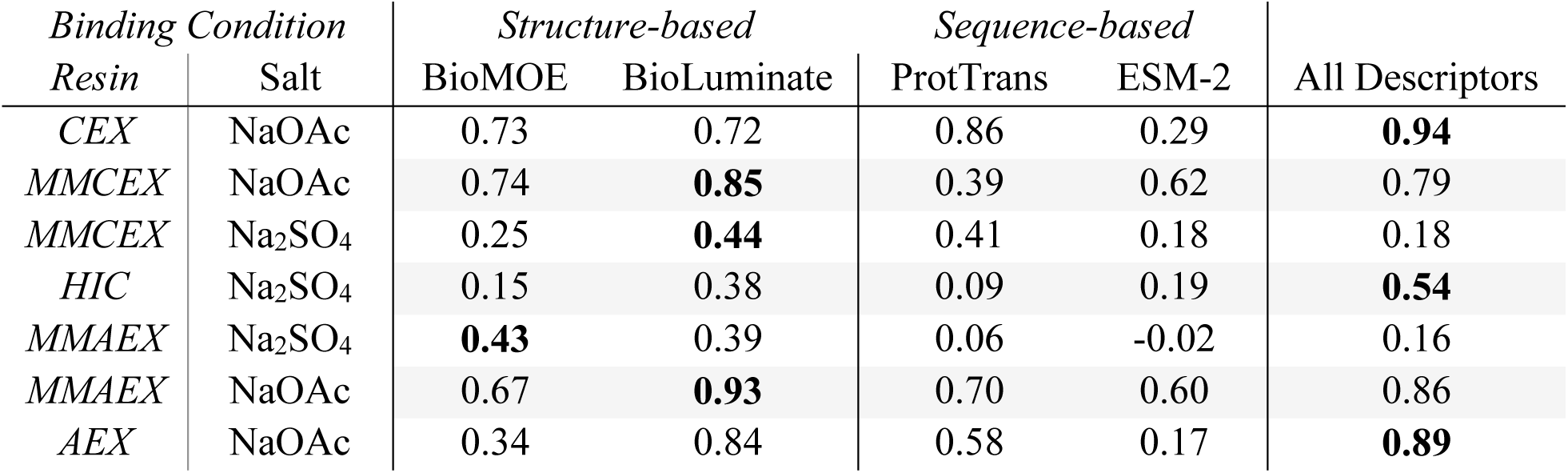
Comparison of withheld test set accuracy (Q^2^) by binding condition and descriptor type. The best models (highest Q^2^) for each binding condition tested are in bold. Training and test set prediction plots and accuracies for the best models are found in Supplemental Figure S5. Selected hyperparameters for the best models are found in Supplemental Table S5. Selected features and features importance are discussed in section 3.5 and Supplemental Figure S8.

| Binding Condition |  | Structure-based |  | Sequence-based |  | All Descriptors |
| --- | --- | --- | --- | --- | --- | --- |
| Resin | Salt | BioMOE | BioLuminate | ProtTrans | ESM-2 |  |
| CEX | NaOAc | 0.73 | 0.72 | 0.86 | 0.29 | **0.94** |
| MMCEX | NaOAc | 0.74 | **0.85** | 0.39 | 0.62 | 0.79 |
| MMCEX | Na<sub>2</sub>SO<sub>4</sub> | 0.25 | **0.44** | 0.41 | 0.18 | 0.18 |
| HIC | Na<sub>2</sub>SO<sub>4</sub> | 0.15 | 0.38 | 0.09 | 0.19 | **0.54** |
| MMAEX | Na<sub>2</sub>SO<sub>4</sub> | **0.43** | 0.39 | 0.06 | -0.02 | 0.16 |
| MMAEX | NaOAc | 0.67 | **0.93** | 0.70 | 0.60 | 0.86 |
| AEX | NaOAc | 0.34 | 0.84 | 0.58 | 0.17 | **0.89** |

Models trained with BioLuminate descriptors are the most predictive in three out of seven binding conditions and second best in three out of seven of the binding conditions evaluated. For three conditions, CEX and AEX in NaOAc and HIC in Na_2_SO_4_, model accuracy improved by the combination of all descriptors. Notably, for CEX in NaOAc, sequence-based ProtTrans descriptors were more predictive than either of the structure-based descriptor sets alone. Models for resins with a predominately charge-based binding mechanism (AEX, CEX, and MMAEX/MMCEX in NaOAc) all achieve test set Q^2^ values above 0.85. However, models for resins with a predominately hydrophobicity-based binding mechanism (HIC/MMAEX/MMCEX in Na_2_SO_4,_ a kosmotrope and promotor of hydrophobic interactions^60^) have Q^2^ scores above 0.43. The observation that for resins in Na_2_SO_4_ test set Q^2^ is lower than training set R^2^ suggests that these models may be overfitting despite optimization of feature selection and KRR regularization. Poor test set generalization may suggest non-optimal side-chain packing in the homology modeling process^11^, that the current set of descriptors may not be capturing some fundamental physical property, or that more data is needed. The lower accuracy of HIC QSPR models aligns with prior studies^11,14,27^ observing poor correlation between structural descriptors and HIC retention time for mAbs (as discussed in section 1.2), and demonstrates the need for development of descriptors which better capture the binding mechanism of hydrophobic interaction chromatography. Despite the lower model accuracy for hydrophobic modes of chromatography, HIC QSPR model predictions are significantly better than random (Q^2^ of zero) and thus can still provide valuable information for assessment of relative risk of molecules in development.

### 2.4 Model Application

Once trained, models can be used to make prospective predictions of purification process fit for molecules in development. PC1 values may be used to compare predicted chromatographic binding behavior to that of molecules in the training set, as shown in Figure 5.

**Figure 5.**
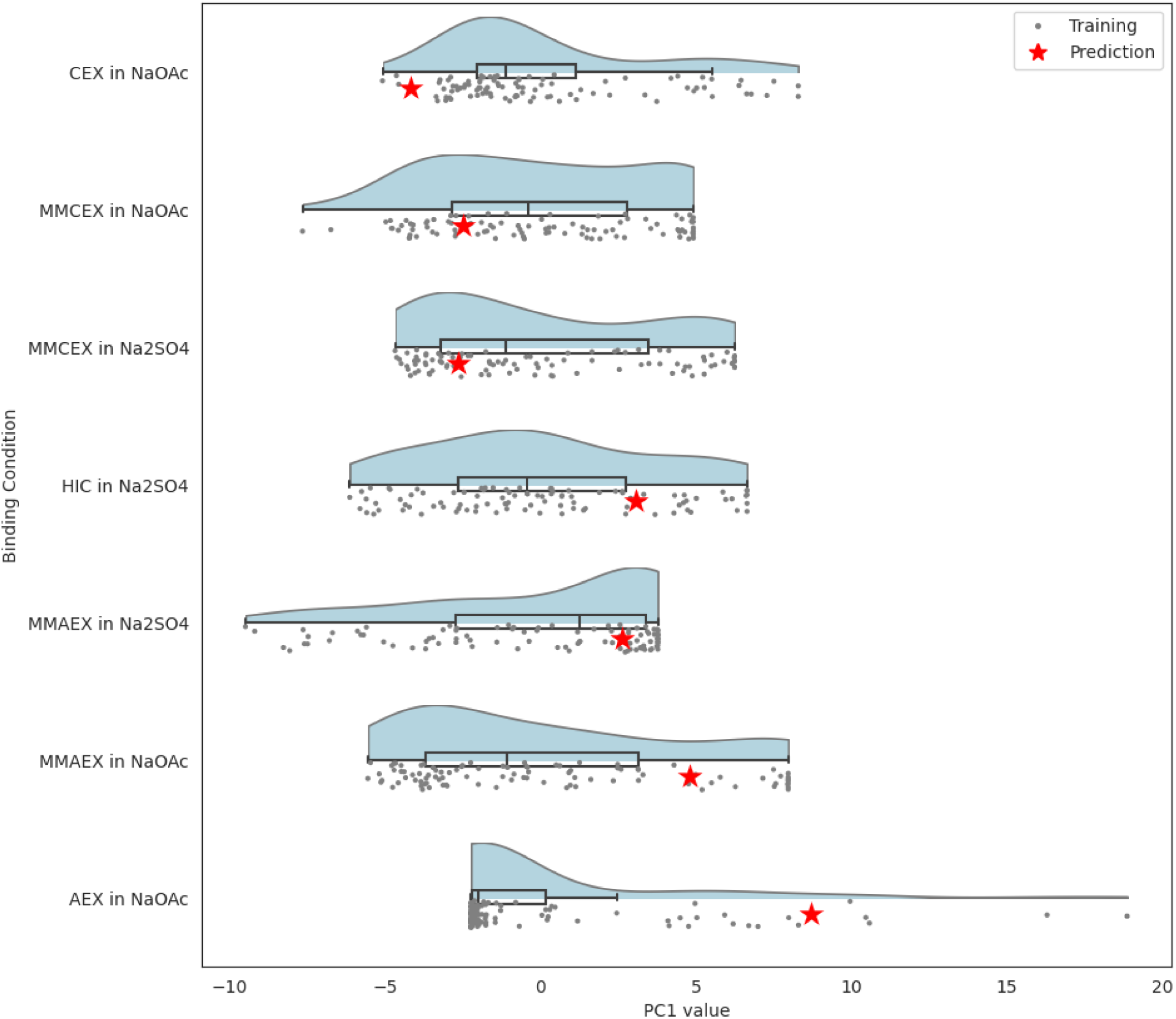
Raincloud plot visualization of predicted molecule PC1 values vs training set PC1 distribution. Raincloud plots combine three elements: a one-dimensional scatter plot visualizing individual molecules, a boxplot visualizing robust distribution statistics (median and interquartile range), and kernel density estimate plot visualizing the distribution of molecules.

At the onset of bioprocess development, the raincloud plot^73^ (Figure 5) can be used to visualize how predicted binding for a new mAb compares to the distribution of prior mAbs. PC1 values within the interquartile range for a given resin indicate typical biophysical properties and binding affinities and thus likely good platform purification process fit, while atypical molecules outside of the interquartile range are flagged as high risk. For example, the predicted molecule shown in Figure 5 demonstrates atypically strong binding to AEX in NaOAc, MMAEX in NaOAc, and HIC in Na_2_SO_4_, and atypically weak binding to CEX in NaOAc. Taken together, these predictions indicate a risk of poor purification process fit for a majority of binding conditions, which is likely to result in challenges during purification process development. Atypically weak binding molecules may face potential risks to facility fit due to low dynamic binding capacity. Atypically strong binding molecules may have potential risk of low yields, challenges with chromatography resin reuse, or necessity of atypically high elution buffer salt concentrations or extreme elution pH, which may increase risk of molecular aggregation or corrosion in stainless steel infrastructure (i.e. high pitting resistance equivalent numbers). Depending on the resin, other risks to consider can include potential to impact to viscosity and column pressure, viral clearance, and separation of other impurities.

Contour plots for the predicted molecule in Figure 5, generated by inverse PCA transform of the predicted PC1 values, are shown in Figure 6.

**Figure 6.**
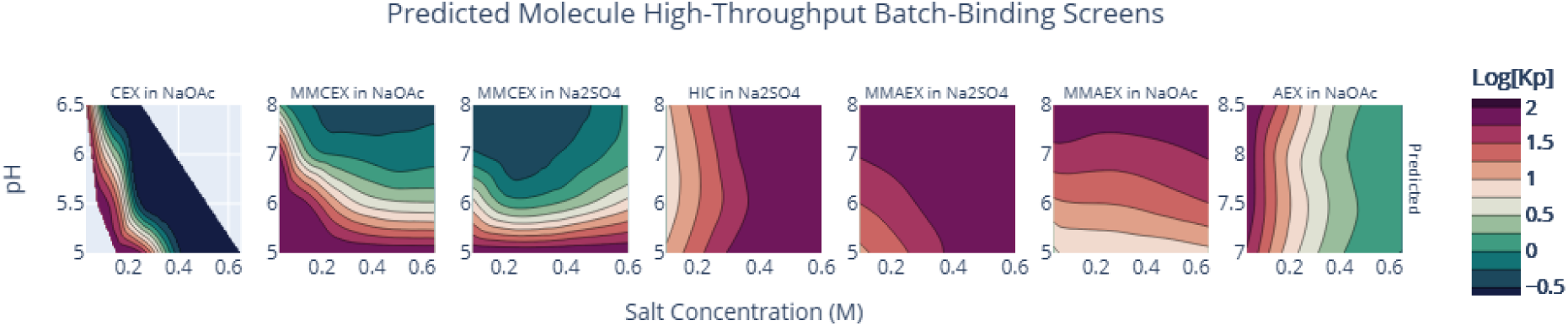
Contour plot visualization of predicted molecule high-throughput batch-binding screen results.

Prediction of PC1 values during purification process fit assessment has multiple benefits including general applicability of PC1 values across various modes of chromatography, and ease of translation between the one-dimensional PC1 representation and the multi-dimensional Log(Kp) contour plot representation. While one-dimensional representations (Figure 5) are well-suited for identification of similar nearest neighbor molecules, contour plots (Figure 6) may be preferred by purification scientists familiar with interpretation of high-throughput batch-binding screen results. These figures can be used in combination when making process development decisions such as selection of resins and process conditions for initial process development experiments. Qualitative assessment of the high-throughput batch-binding screen contour plots allows researchers to identify potential operating conditions, such as regions with low Log(Kp) values with potential to operate in flow-through mode, and regions of transition from high to low Log(Kp) values with potential to operate in bind and elute mode.

The presented QSPR modeling strategy can also be used to identify similar nearest neighbor molecules. Historical knowledge from nearest neighbor molecules with similar PC1 values may provide a useful starting point for identifying potential molecular liabilities, and informing process development decisions such as selection of resins and process conditions. Figure 7 provides an illustration of the utility of predicted PC1 values for identification of similar nearest neighbor molecules to inform process development for CEX in NaOAc. Similar figures for every combination of resin and salt type may be found in Supplemental Figure S5.

**Figure 7.**
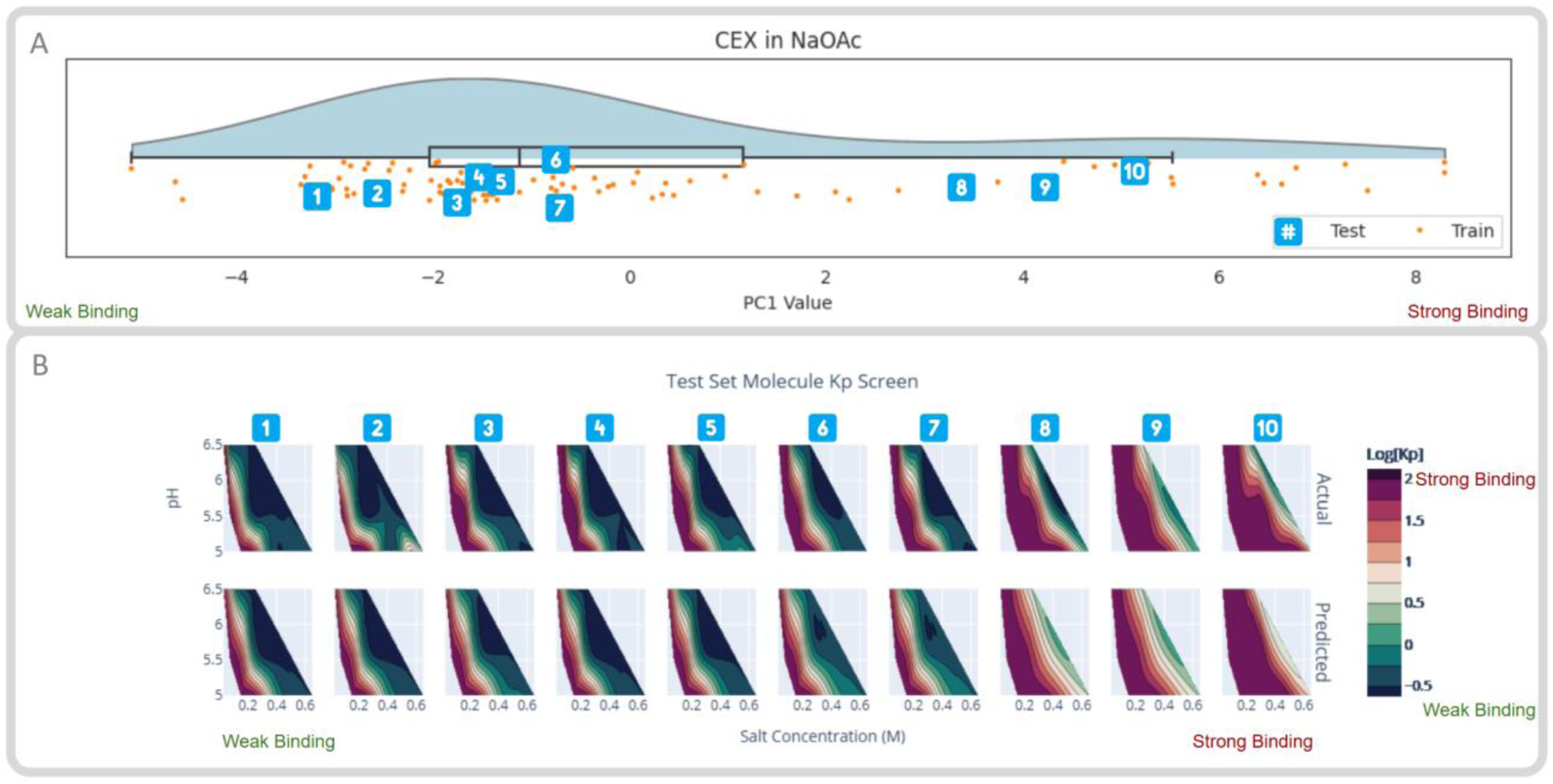
Model prediction outputs for withheld test set molecules. (A) illustrates the distribution of predicted test set molecule PC1 values on the raincloud plot. (B) visualizes the similarity of Log(Kp) contour plots with similar PC1 values for CEX in NaOAc using test set molecules. Molecule 5 was removed when calculating the test set metrics due to high sequence similarity to training set molecules (see section 2.4, Supplemental Figure S4) Comparison of the rows demonstrates the similarity of the actual Log(Kp) values to predicted Log(Kp) values derived from inverse PCA transform of the predicted PC1 value.

As previously discussed, raincloud plots such as Figure 7A are useful for identification of similar molecules in order to leverage historical knowledge. In Figure 7B, test molecules with similar predicted PC1 values have qualitatively similar actual Log(Kp) contour plots. Therefore, although information may be lost due to imperfect PCA compression or QSPR model prediction, the proposed modeling strategy still demonstrates utility for prediction of chromatographic binding behavior and identification of nearest neighbor molecules whose actual experimental results are similar.

While the experimental dataset in the present work limits model predictions to standard mAbs, comparison of relative binding for Fabs or individual arms of multi-specific antibodies might be accomplished by making predictions for Fabs on a common Fc framework. Prior work from Saleh, Hess, *et al.* demonstrated good results^32,35,36^ for models trained on diverse molecular formats including Fabs, bispecific, and multi-specific antibodies. Addition of training data for other molecular formats to the present QSPSR dataset could also potentially extend utility of the models presented in this work.

### 2.5 Analysis of Model Feature Importance

QSPR models are typically trained with high-dimensional data with many more descriptors than training datapoints. The abundance of descriptors arises from calculating descriptors in multiple ways (at multiple pH levels, with multiple hydrophobicity scales, weighted by various criteria for solvent accessibility and surface patchiness, globally and for multiple protein subdomains). As such, QSPR models must deal with the pervasive challenges of non-predictive features and redundant features (i.e. multicollinearity, illustrated in Supplemental Figure S7). Consequently, QSPR models depend on feature selection, and analysis of model feature selection and feature importance can reveal the basis of model predictions and inform hypotheses about mechanisms of chromatographic binding. For the models with the highest Q^2^ value (bold values in Table 2), the type and number of descriptors selected by RFE is shown in Figure 8.

**Figure 8.**
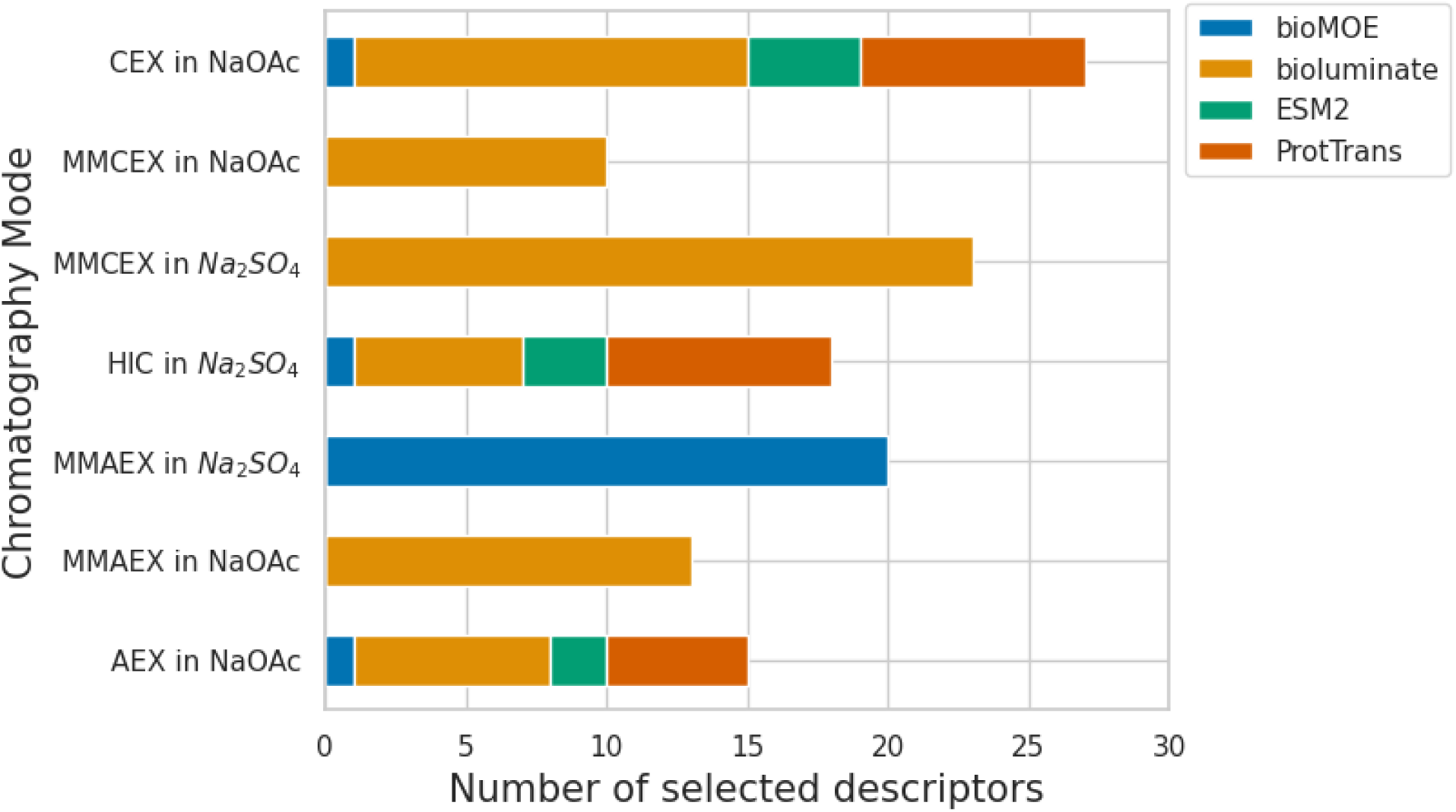
Type and quantity of features selected by model-based recursive feature elimination. Feature importance plots, illustrating the features selected for each model, are provided in Supplemental Figure S8.

In all cases, RFE selects fewer than 30 features in the final model, below the upper limit of the Bayesian Optimization search range (Supplemental Table S4), illustrating the importance of RFE to prevent model overfitting. Higher number of features for Na_2_SO_4_ models, coupled with decreased test set performance (Q^2^ lower than R2), indicates the model pipeline may be overfitting on these data sets. More data or more extensive hyperparameter tuning could help improve these models in the future. In most cases, BioLuminate descriptors make up the plurality of descriptors selected. However, examination of descriptor counts agnostic of relative feature importance can be misleading.

For analysis of the relative impact of features on model outputs, discussion of SHAP^74^ feature importance values is found in Supplemental Figure S8. In general, charge descriptors (charge patch, dipole moment, and zeta potential) have the highest feature importance for ion exchange (AEX/CEX) and mixed mode (MMAEX/MMCEX) resins in NaOAc. This suggests that for the conditions tested, mixed mode resins are primarily binding by the ion exchange mechanism. In alignment with most other resins in NaOAc, BioLuminate descriptors are the most important for CEX in NaOAc (8 of the 10 highest SHAP scores). For AEX in NaOAc, half of the highest SHAP scores for the best model were protein language model descriptors. However, the combination of all descriptors only improved model accuracy by 0.05 in comparison to BioLuminate descriptors alone (Table 2). By contrast, predictions for HIC in Na_2_SO_4_ primarily rely on protein language model descriptors (8 of the 10 highest SHAP scores) and the combination of all descriptors improved model accuracy by 0.16 over BioLuminate descriptors alone (Table 2). This recapitulates the need for improved structural descriptors for hydrophobicity and highlights the potential of protein language models as a complementary molecular descriptor generation strategy. Compared to models in NaOAc, MMAEX and MMCEX in Na_2_SO_4_ display higher feature importance for aromatic groups and hydrogen bond acceptors respectively.

## 3 Discussion

In conclusion, this study introduces a novel modeling strategy for *in silico* prediction of chromatographic binding behavior across wide ranges of pH and salt concentration, in support of purification process fit assessment for mAbs. This study also proposes application of PCA to compress multi-dimensional data from high-throughput batch-binding screens into a one-dimensional metric for comparison of molecular chromatographic binding affinity.

The modeling strategy benchmarks the performance of structural-based molecular descriptors and sequence-based protein language model embeddings across five different chromatography resins in two different salt backgrounds. In order to prevent overfitting on high-dimensionality data, the modeling strategy uses a multi-stage feature selection strategy leveraging a correlation-based feature filter, model-based recursive feature selection, and a regularized kernel ridge regressor. Applying this modeling strategy to prediction of PC1 values achieves test set Q^2^ values over 0.85 for models of ion exchange and multimodal chromatography in NaOAc, and over 0.4 for models of multimodal and hydrophobic interaction chromatography in Na_2_SO_4_.

Notably, our findings underscore the practical effectiveness of *in silico* QSPR predictions as viable alternatives to experimental high-throughput batch-binding screens for assessment of chromatographic binding behavior and downstream bioprocess fit. Our models support identification of molecules with atypical chromatographic binding behavior, identification of nearest neighbor molecules with similar binding affinities supporting better use of historical knowledge, and ultimately offer insights to the selection of appropriate resins and purification process conditions during purification process development. This research represents another step towards streamlining the early-stage development of mAbs, and mitigating timeline pressures and material constraints through the power of predictive modeling.

While these models help inform risk assessment, further improvements to model accuracy are needed. Many current QSPR model applications for chromatography focus on qualitative assessment of molecular binding affinity (i.e. categorical flags for purification process development risks, ordinal ranking of relative binding affinity in order to identify nearest neighbor molecules) because of (1) small data sizes and (2) imperfect biophysical molecular descriptors. Future work should look at expanded data set sizes and new approaches for descriptors that may be more physically relevant to chromatography binding mechanisms. To reduce risk of overfitting on poorly predictive HIC descriptors, improved descriptors are needed. One possible area of research is development of structure-based biophysical descriptors better capturing mechanisms of hydrophobic binding. Use of mAb-specific deep learning models for structure prediction and sequence embedding are promising potential areas for evaluation in future work. Use of alternative machine learning regression algorithms and transfer learning with larger datasets are also potentially interesting topics of future work. To reduce the risk of model over-interpretation, future work may consider prediction uncertainty. Improved QSPR model accuracy could expand model use to quantitative predictions, with the potential to replace experiments, debottleneck selection of purification process operating conditions, resulting in more robust purification processes, accelerated purification process development, and more rapid development of mAbs at lower cost to patients and society.

## 5 Abbreviations

AEX: anion exchange chromatography
CDR: complimentary-determining region
CEX: cation exchange chromatography
Cryo-EM: cryogenic electron microscopy
CV: cross validation
Fab: fragment antigen-binding region
Fv: variable fragment domain
GaMD: gaussian accelerated molecular dynamics
HIC: hydrophobic interaction chromatography
Ig: immunoglobulin
Kp: partition coefficient
KRR: kernel ridge regression
K-SVR: kernel support vector regression
mAb: monoclonal antibody
MAE: mean absolute error
MMAEX: multimodal anion exchange chromatography
MMCEX: multimodal cation exchange chromatography
MOE: molecular operating environment
Na_2_SO_4_: sodium sulfate
NaOAc: sodium acetate
PC1: first principal component
PCA: principal component analysis
PDB: protein data bank
pI: isoelectric point
PLS: partial least squares
_Q_^2^: test set coefficient of determination
QSPR: quantitative structure-property relationship
_R_^2^: training set coefficient of determination
RFE: recursive feature elimination
RR: ridge regression
SAP: spatial aggregation propensity
SMA: steric mass action

## 6 Author Contributions

Conceptualization: Andrew Maier, Minjeong Cha, Sean Burgess, Ambrose Williams.

Design & Execution of Experiments: Rituparna Sengupta, Liliana Yee, Kelly O’Connor, Nicole Ott, Neeraja Sundar Rajan, Josephine Neyyan, Carlos Cuellar, Soo Kim, Ambrose Williams.

Modeling Methodology: Andrew Maier, Minjeong Cha, Sean Burgess, Amy Wang.

## 7 Acknowledgements

The authors would like to acknowledge experimental and scientific support of their colleagues in the Cell Culture and Bioprocess Operations, Purification, Microbiology and Virology, and Pharmaceutical Development departments at Genentech, in particular those named below.

Support of experimentation: Jonathan Zarzar, Juan Lopez, Liliana Yee, Samantha Leung, Bryan Celarbo, Romie Aposta, Minerva Olaya, Andre Hung Phan.

Support of modeling: Connor Thompson, Jessica Lyall, Michael Chinn, Saeed Izadi, Eliott Park, Nandhini Rajogopal, Derek Lee, Benjamin Tran.

## 9 Additional Funding

The author(s) reported that there is no funding associated with the work featured in this article.

