## Supplemental Figures for "Predicting purification process fit of monoclonal antibodies using machine learning"

### Supplementary Information

#### Section A) Feature Correlation to PC1

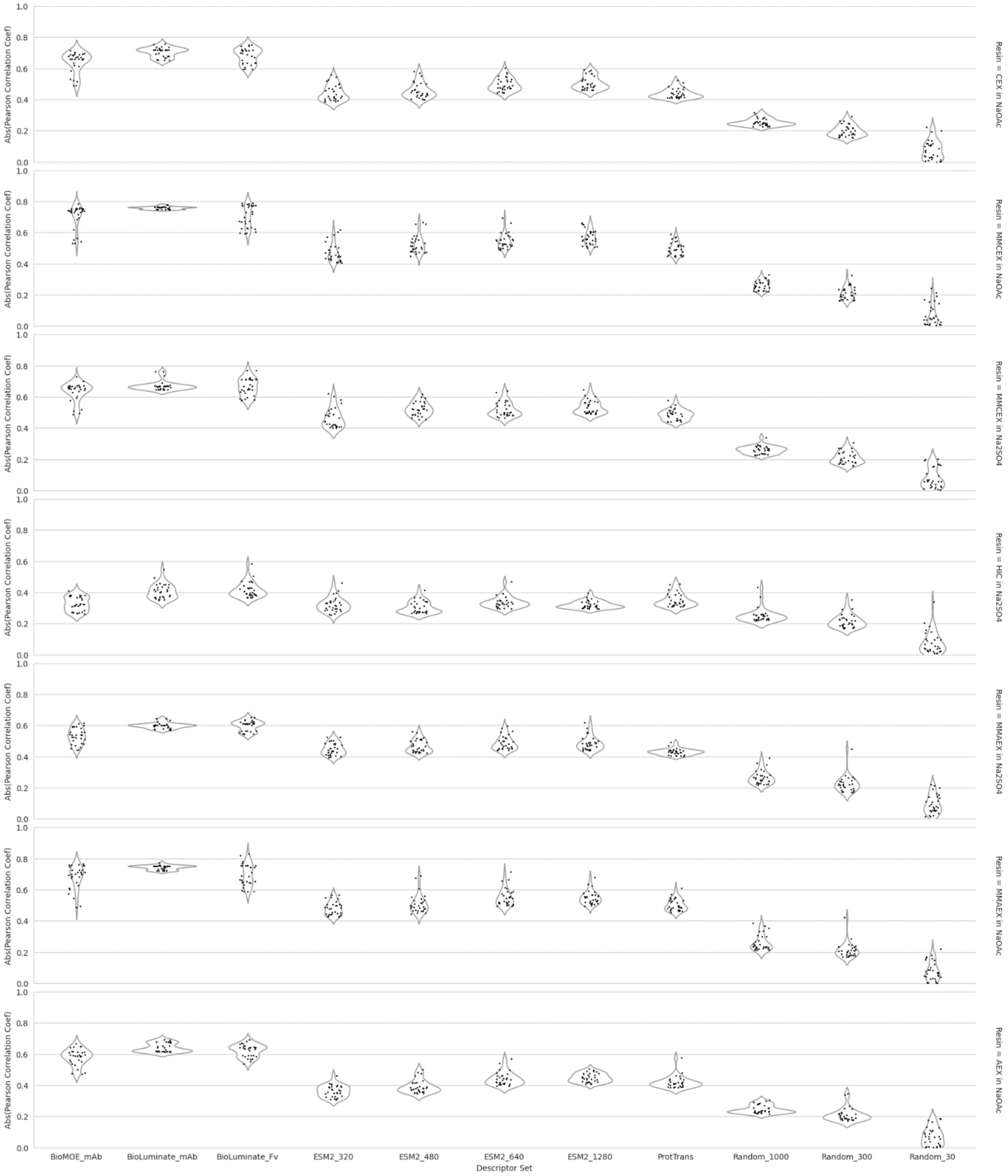


*Figure S1 – Distribution of the 30 most Highly Correlated Descriptors for Pearson Correlation between Descriptors and PC1 Values. Structures used for BioLuminate structural descriptors and the number of descriptors for protein language model and random descriptors are indicated by the x-axis labels.*

In order to qualitatively assess the relevance of different descriptor sets to PC1 prediction, the distribution of Pearson correlation coefficients between descriptors and PC1 values are compared in Figure S1.

Random descriptors sets of various sizes are included as a negative control. Larger descriptor sets may contain more highly correlated descriptors by random chance. The increase in Pearson correlation observed across random descriptor sets results from visualizing only the 30 mostly highly correlated descriptors. All other descriptor sets have significantly higher Pearson correlation than the random descriptor sets.

For BioLuminate descriptors, full-mAb descriptors have higher Pearson correlation than Fv descriptors across all binding conditions. For full-mAb structural descriptors, BioLuminate descriptors have higher Pearson correlation than BioMOE. Given these findings, full-mAb structural descriptors are used for model development.

For sequence-based descriptors, ESM and ProtTrans have similar Pearson correlation coefficients. Larger numbers of sequence-based descriptors correspond to protein language models with more parameters and higher dimensional embeddings. For ESM-2 descriptor sets, higher Pearson correlation coefficients are observed for larger embeddings. However, since the magnitude of increase in Pearson correlation across ESM-2 embedding sizes is similar to the increase of Pearson correlation for across randomly generated descriptor sets, it’s possible that higher correlations observed for larger ESM-2 embeddings may be attributable to the number of descriptors rather than their quality. In order to reduce the risk of overfitting on a high-dimensional embedding with the small QSPR dataset, the smallest ESM-2 model is used for model development.

#### Section B) Sequence Identity Analysis

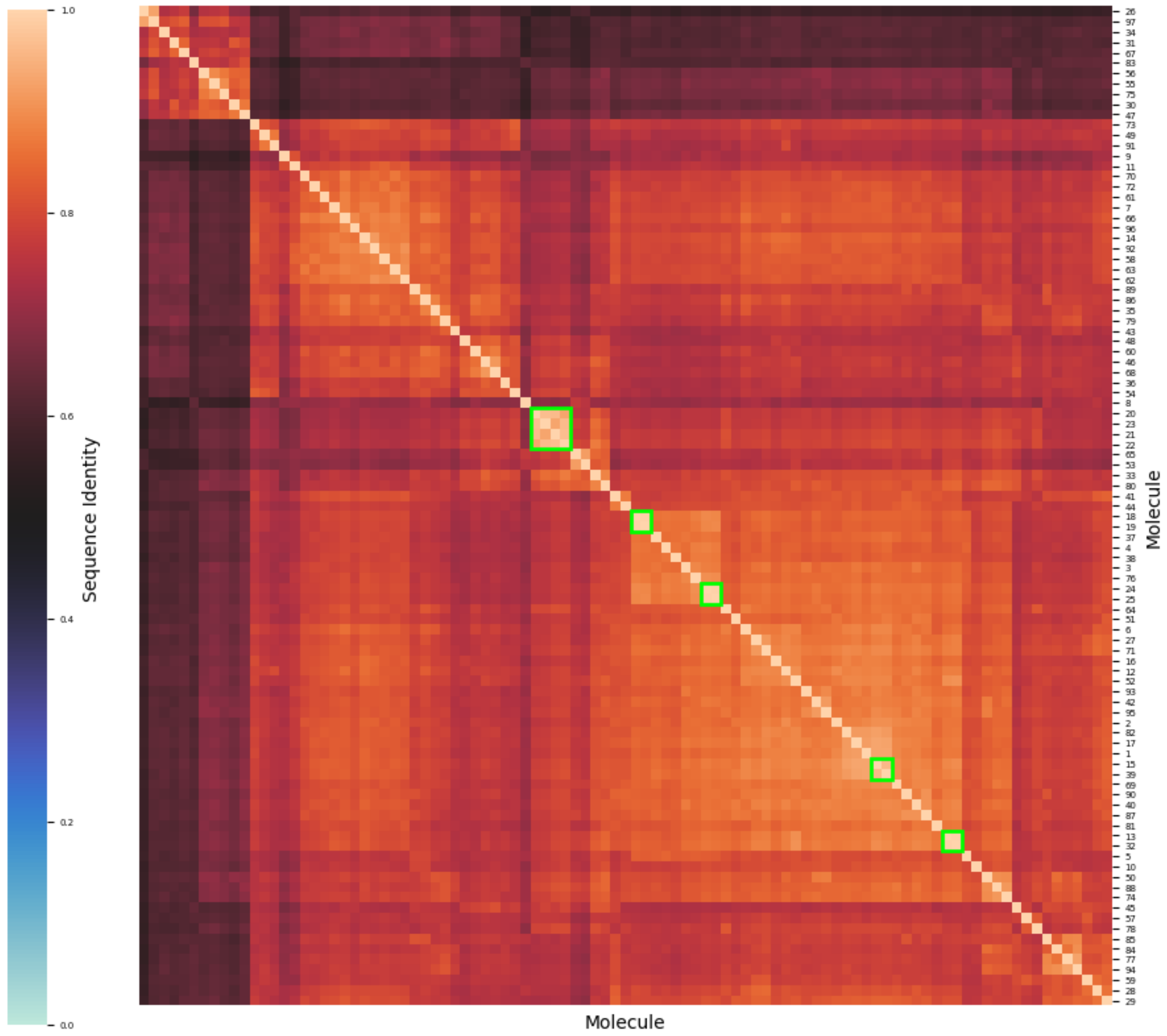


*Figure S2 – mAb Sequence Identity Clustermap. Sequence alignment and Sequence Identity between full mAb sequences in the dataset are performed with Biopython. Clustermap visualization illustrates groups of molecules with sequence identity above 95% (green squares). In total, 5 clusters were identified each with 2-4 members. Additional details of the molecules removed for each binding condition tested are provided in Supplemental Figure S6.*

### Section C) Principal Component Analysis

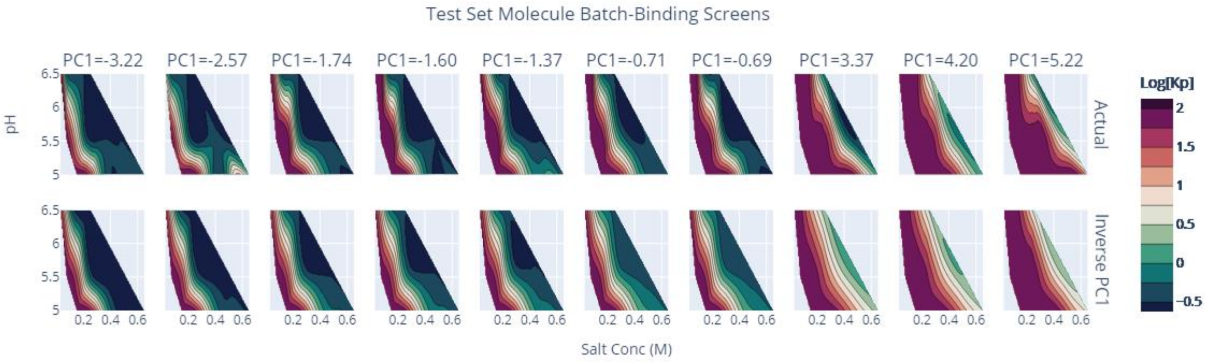


*Figure S3.1 – Comparison of Actual Log Kp values vs Log Kp values from inverse PCA transform of PC1 values for CEX in NaOAc*

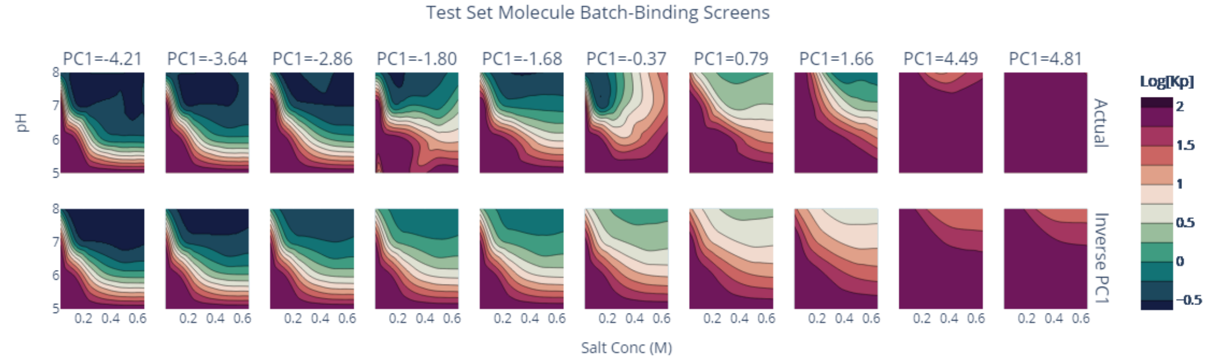


*Figure S3.2 – Comparison of Actual Log Kp values vs Log Kp values from inverse PCA transform of PC1 values for MMCEX in NaOAc*

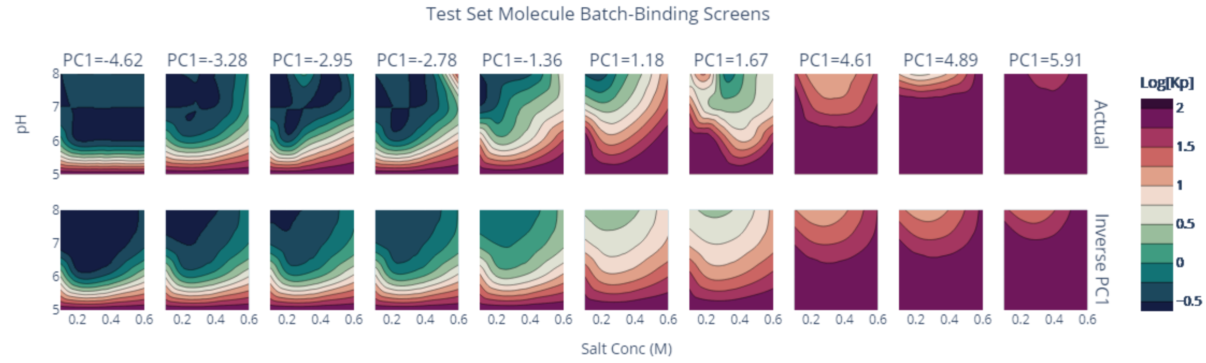


*Figure S3.3 – Comparison of Actual Log Kp values vs Log Kp values from inverse PCA transform of PC1 values for MMCEX in Na2SO4*

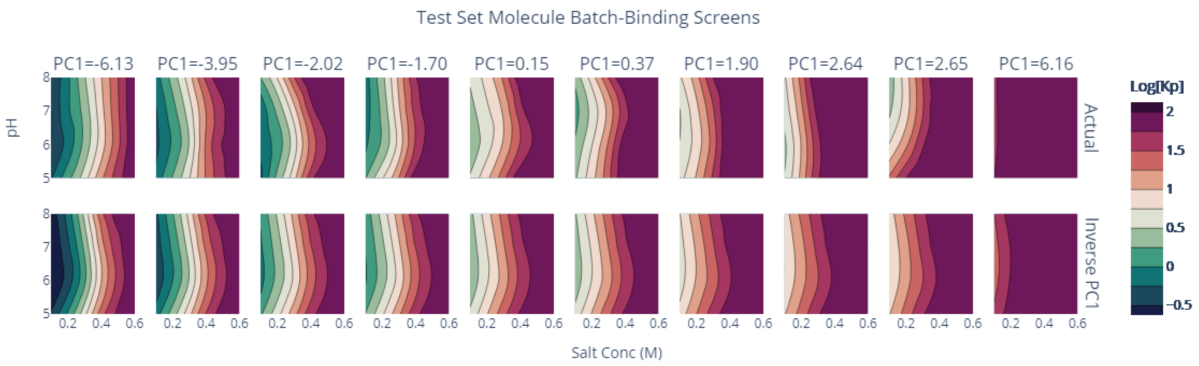


*Figure S3.4 – Comparison of Actual Log Kp values vs Log Kp values from inverse PCA transform of PC1 values for HIC in Na2SO4*

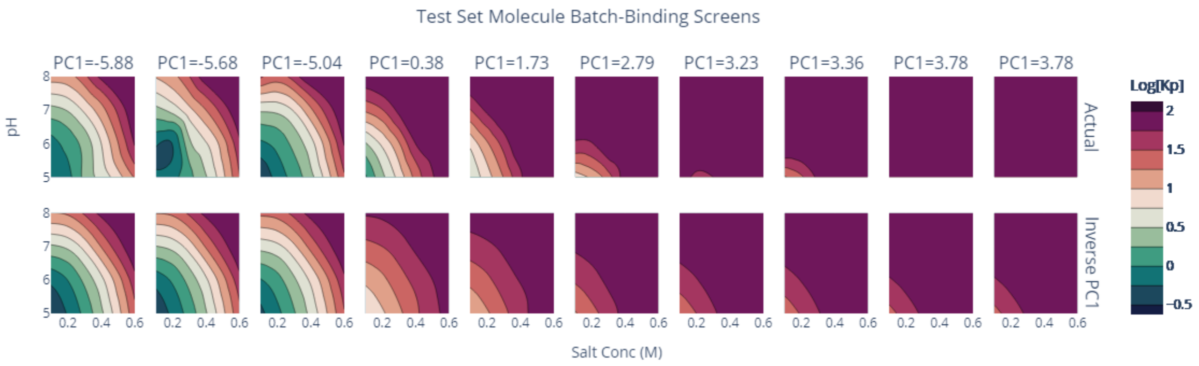


*Figure S3.5 – Comparison of Actual Log Kp values vs Log Kp values from inverse PCA transform of PC1 values for MMAEX in Na2SO4*

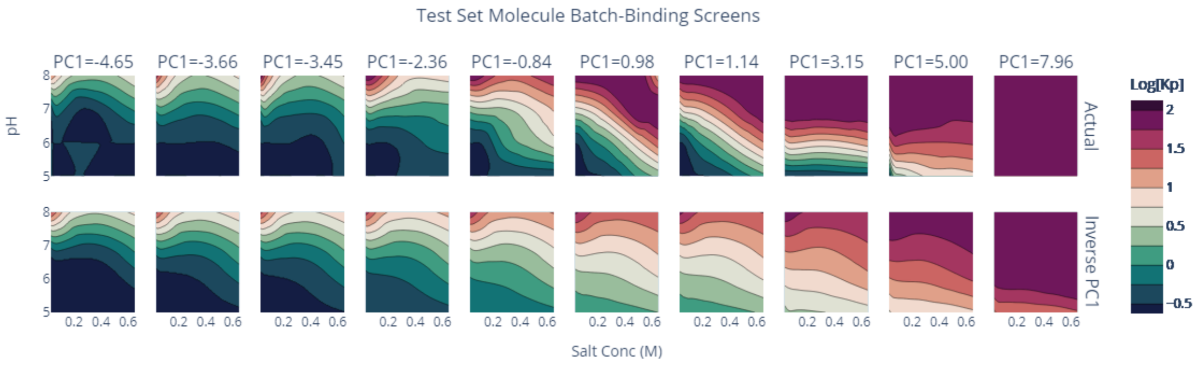


*Figure S3.6 – Comparison of Actual Log Kp values vs Log Kp values from inverse PCA transform of PC1 values for MMAEX in NaOAc*

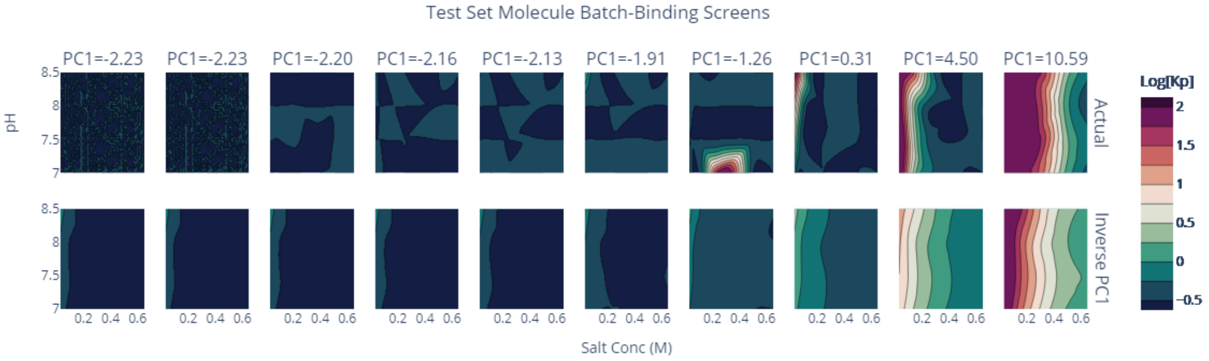


*Figure S3.7 – Comparison of Actual Log Kp values vs Log Kp values from inverse PCA transform of PC1 values for AEX in NaOAc*

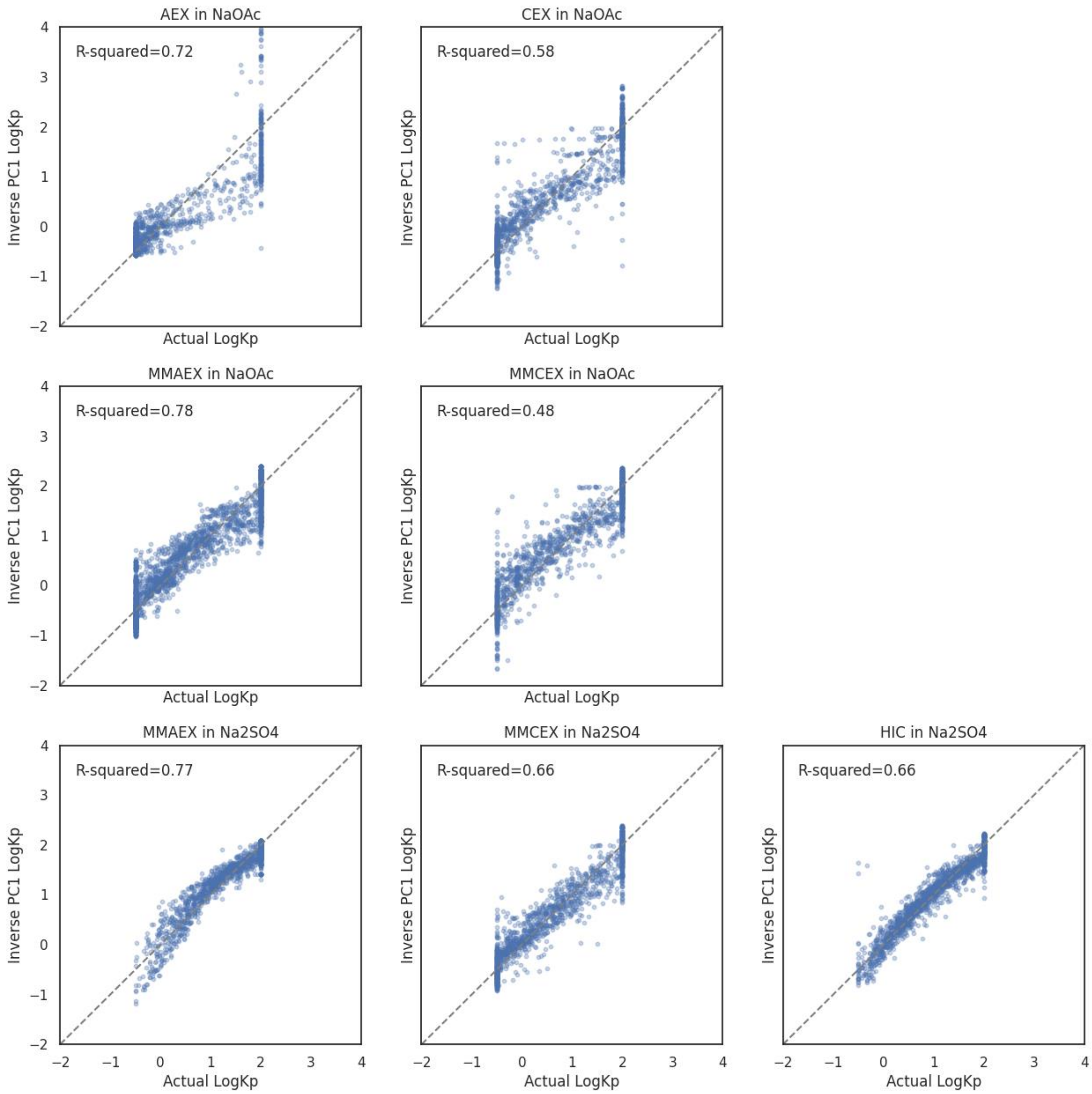


*Figure S4 – Comparison of Actual Log Kp values vs Log Kp values from inverse PCA transform of PC1 values.*

### Section D) Model Accuracy

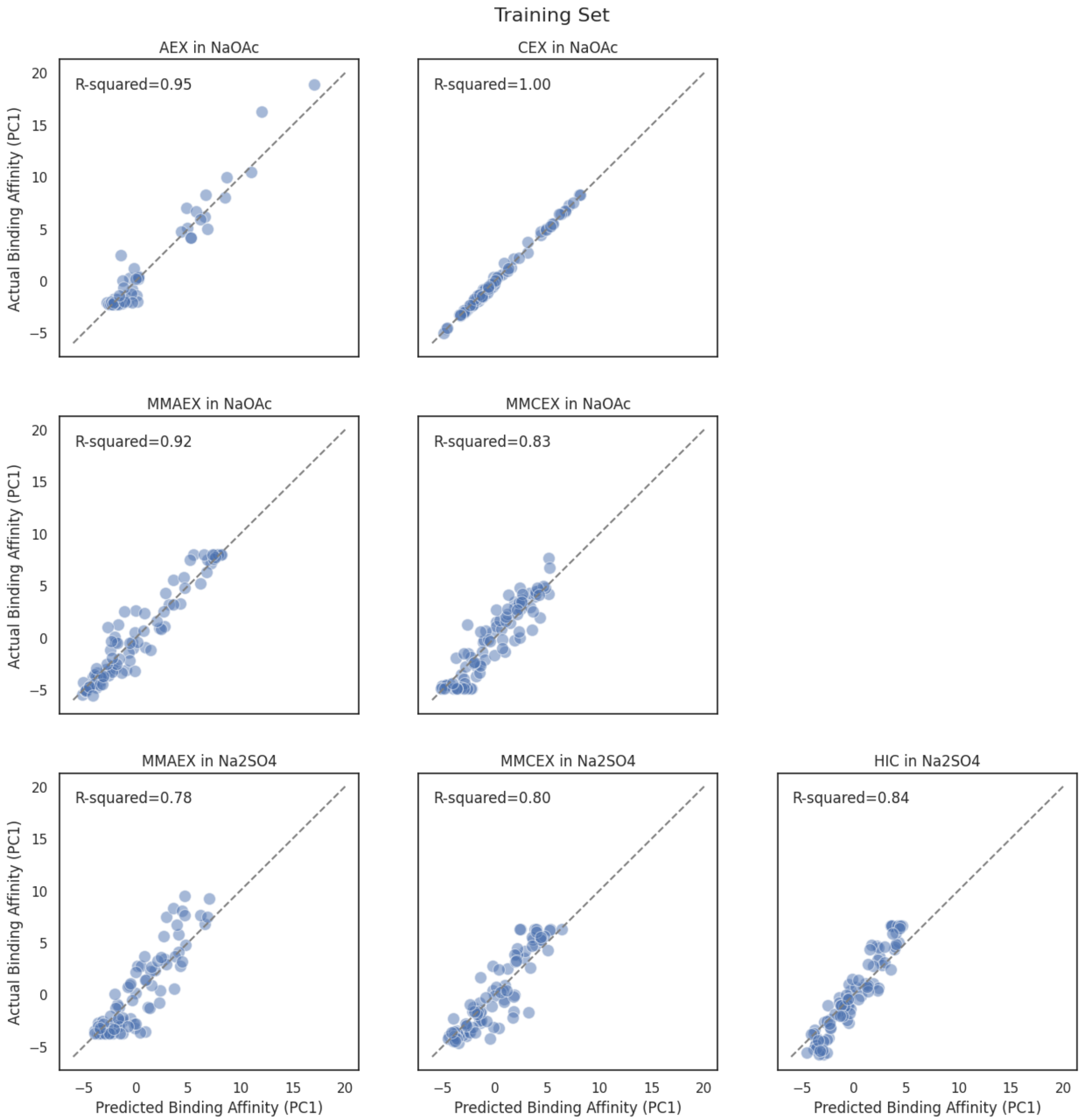


*Figure S5.1 – Prediction Plots for training set molecules. For each binding condition, the model for the descriptor set with the highest test set accuracy is plotted.*

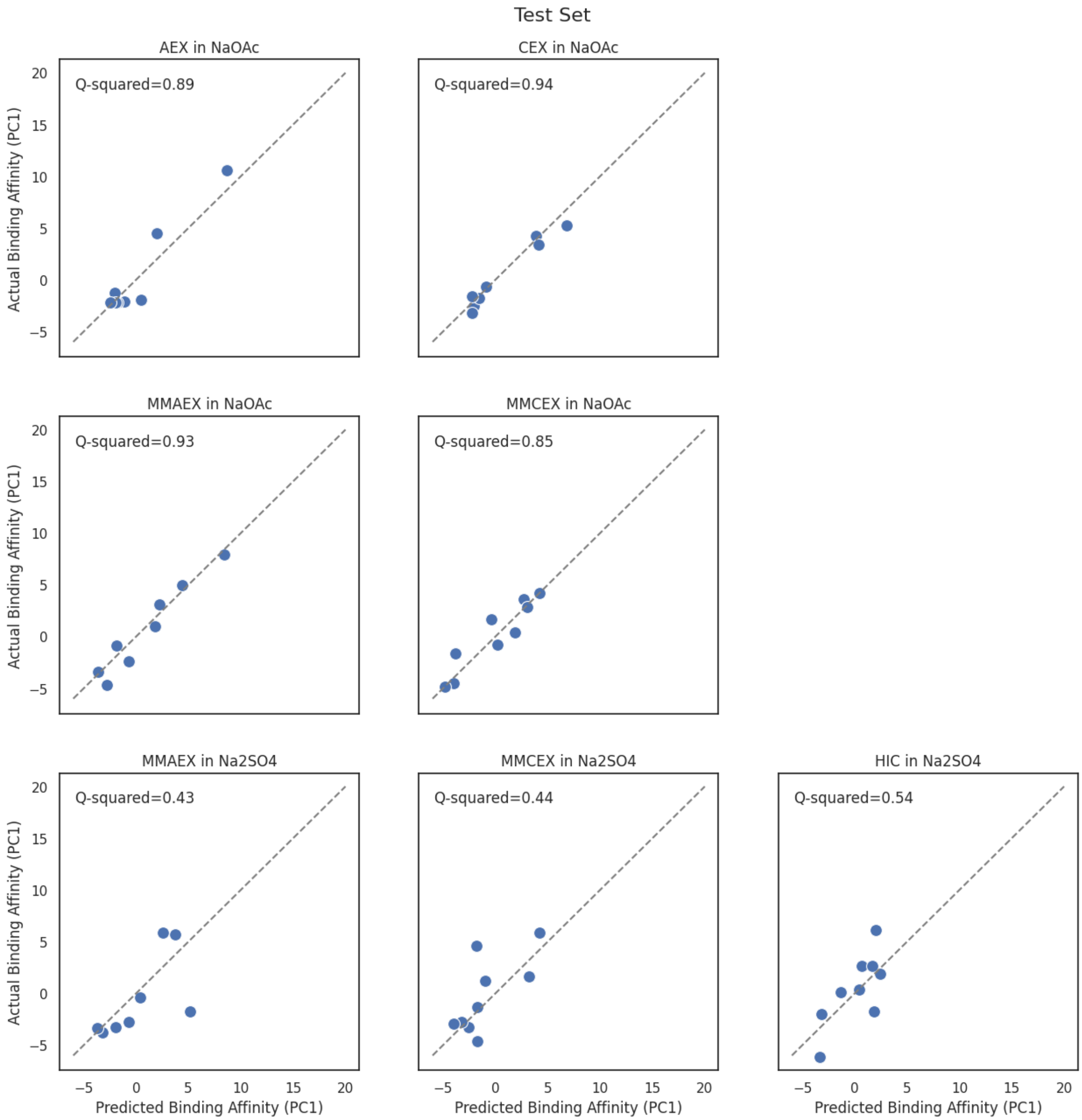


*Figure S5.2 – Prediction Plots for test set molecules. For each binding condition, the model for the descriptor set with the highest test set accuracy is plotted. Molecules with high sequence similarity to training set molecules were excluded when calculating test set metrics (see manuscript section 2.4, Supplemental Figures S4 and S6).*

### Section E) Model Application

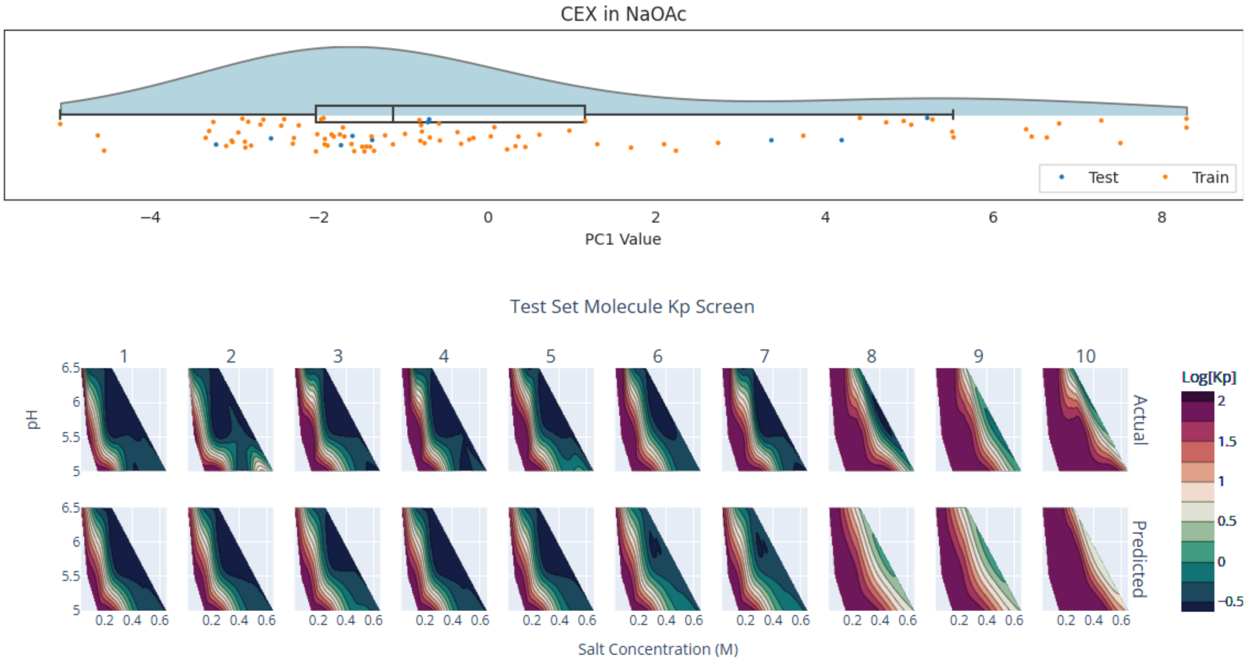


*Figure S6.1 – Cation Exchange Chromatography in Sodium Acetate. Molecule 5 was removed when calculating the test set metrics due to high sequence similarity to training set molecules (see manuscript section 2.4, Supplemental Figure S4).*

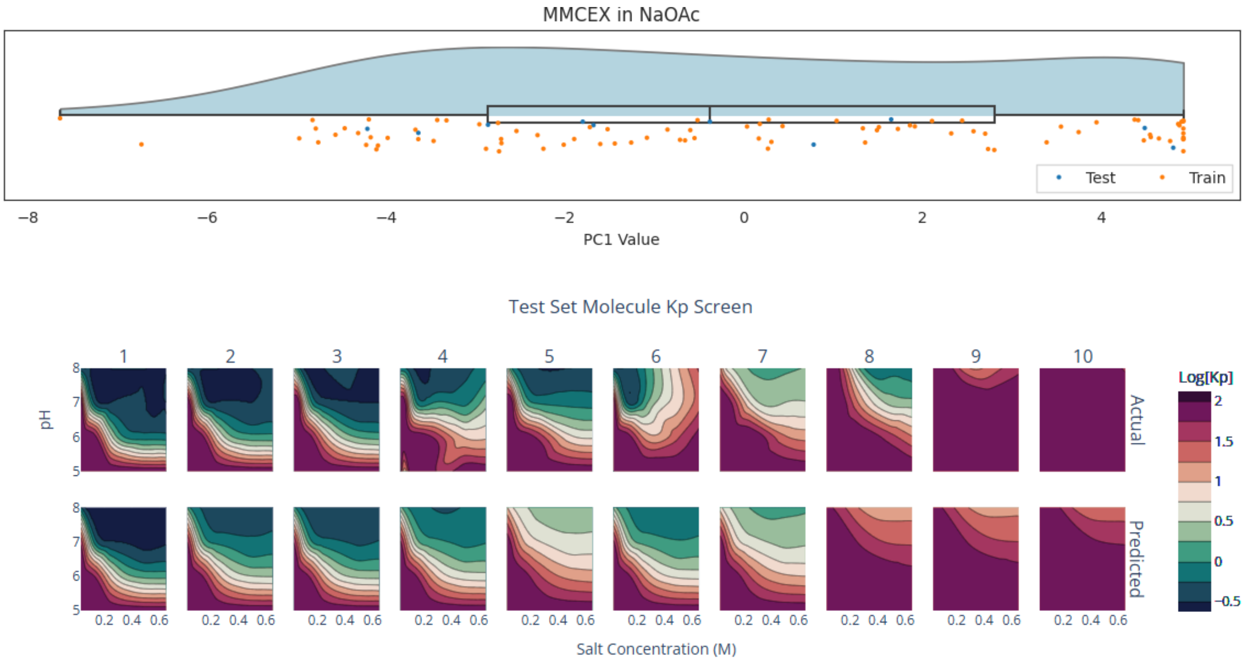


*Figure S6.2 – Mixed Mode Cation Exchange Chromatography in Sodium Acetate. Molecule 4 was removed when calculating the test set metrics due to high sequence similarity to training set molecules (see manuscript section 2.4, Supplemental Figure S4).*

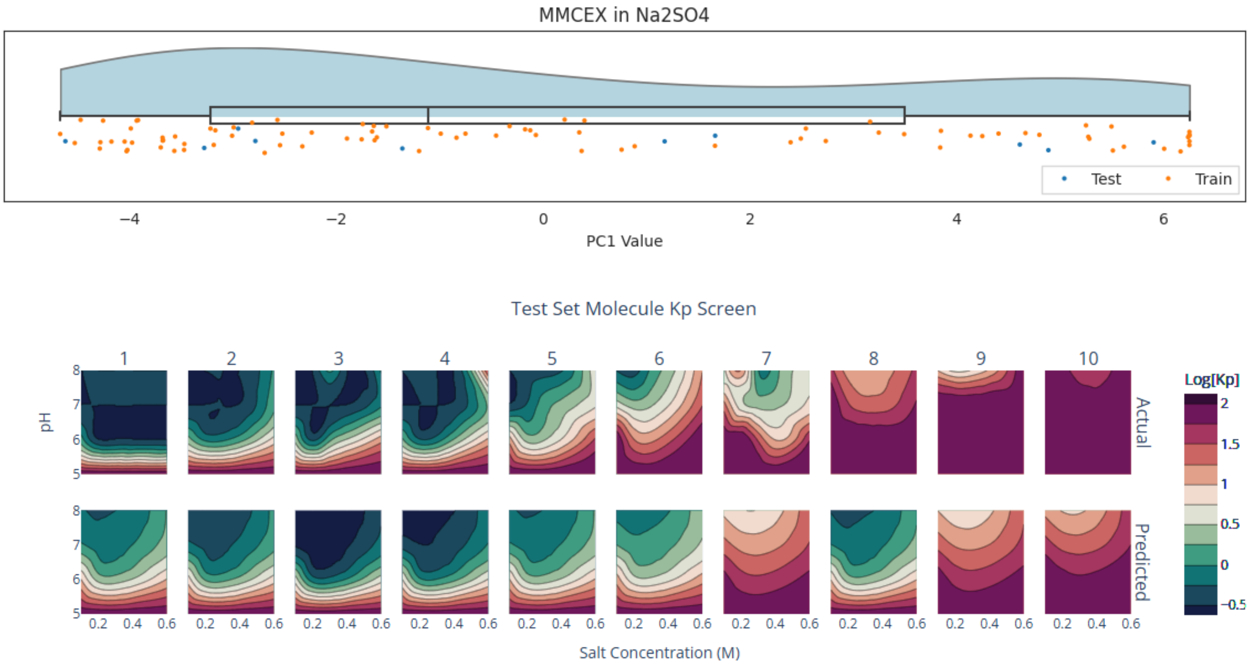


*Figure S6.3 – Mixed Mode Cation Exchange Chromatography in Sodium Sulfate. Molecule 9 was removed when calculating the test set metrics due to high sequence similarity to training set molecules (see manuscript section 2.4, Supplemental Figure S4).*

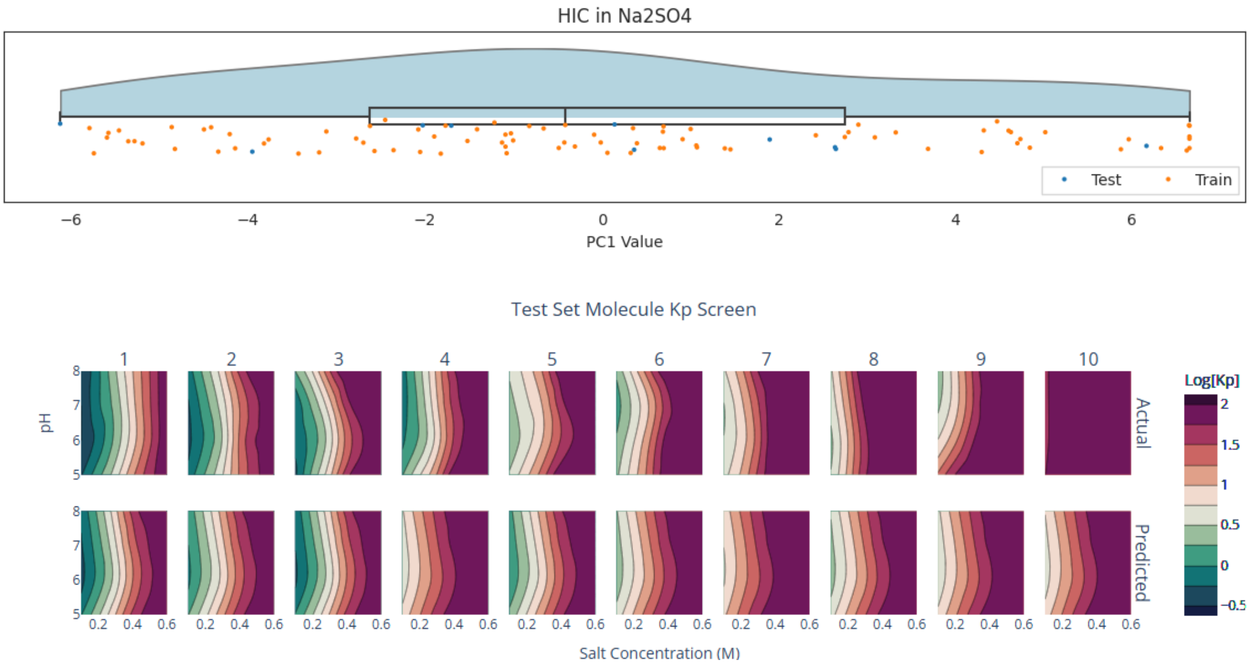


*Figure S6.4 – Hydrophobic Interaction Chromatography in Sodium Sulfate. Molecule 2 was removed when calculating the test set metrics due to high sequence similarity to training set molecules (see manuscript section 2.4, Supplemental Figure S4).*

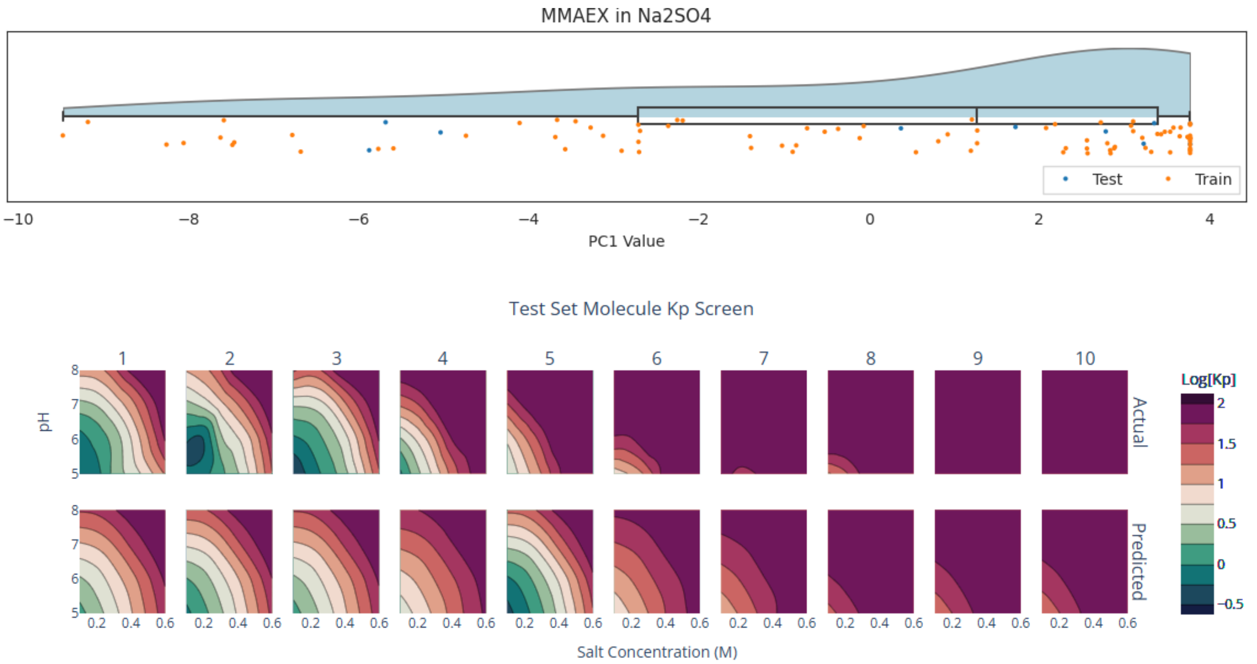


*Figure S6.5 – Mixed Mode Anion Exchange Chromatography in Sodium Sulfate. Molecule 3 was removed when calculating the test set metrics due to high sequence similarity to training set molecules (see manuscript section 2.4, Supplemental Figure S4).*

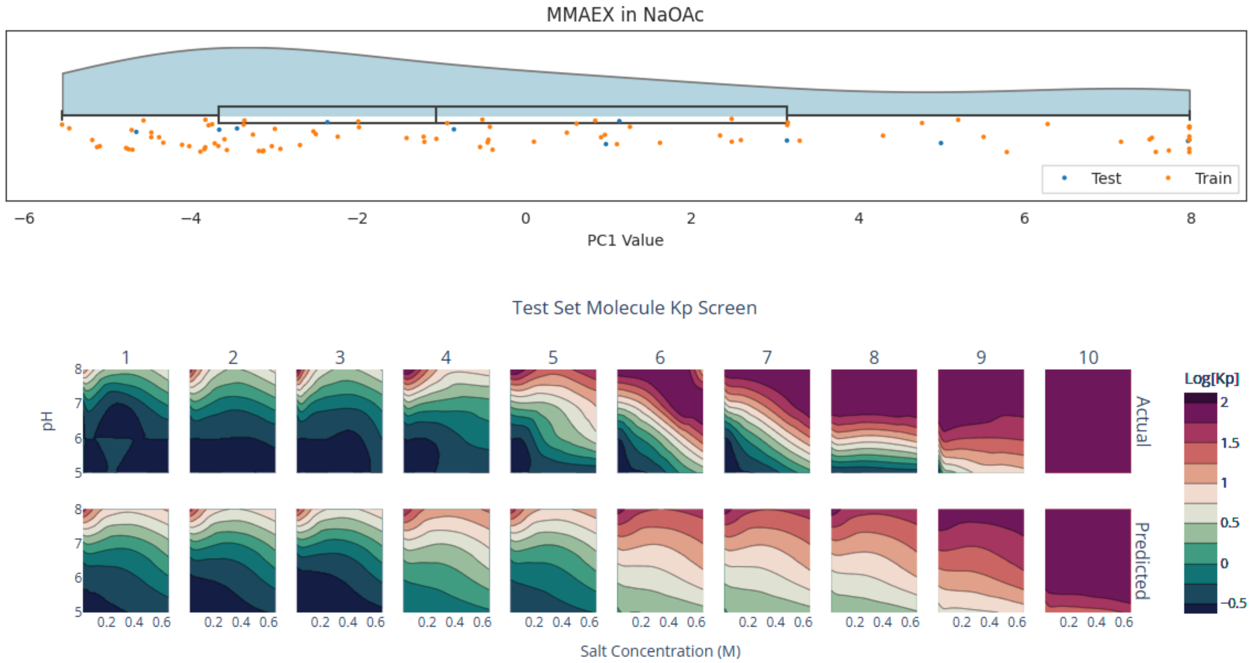


*Figure S6.6 – Mixed Mode Anion Exchange Chromatography in Sodium Acetate. Molecules 2 and 7 were removed when calculating the test set metrics due to high sequence similarity to training set molecules (see manuscript section 2.4, Supplemental Figure S4).*

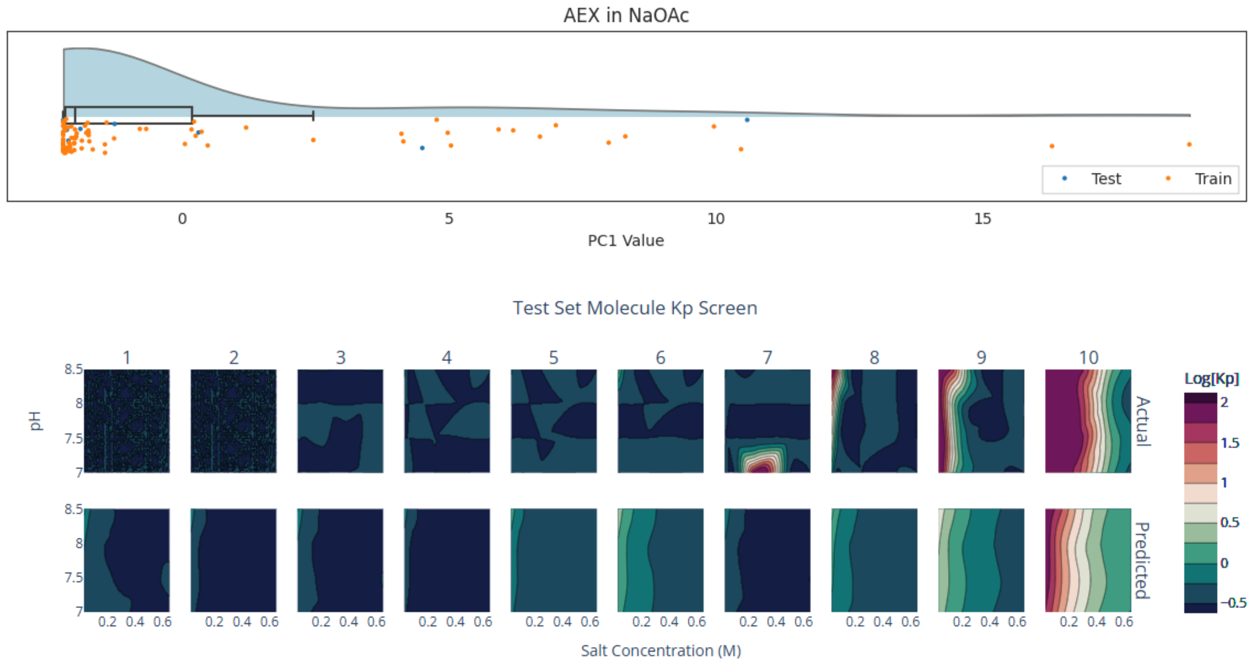


*Figure S6.7 – Anion Exchange Chromatography in Sodium Acetate. Molecule 8 was removed when calculating the test set metrics due to high sequence similarity to training set molecules (see manuscript section 2.4, Supplemental Figure S4).*

#### Section F) Feature Multicollinearity

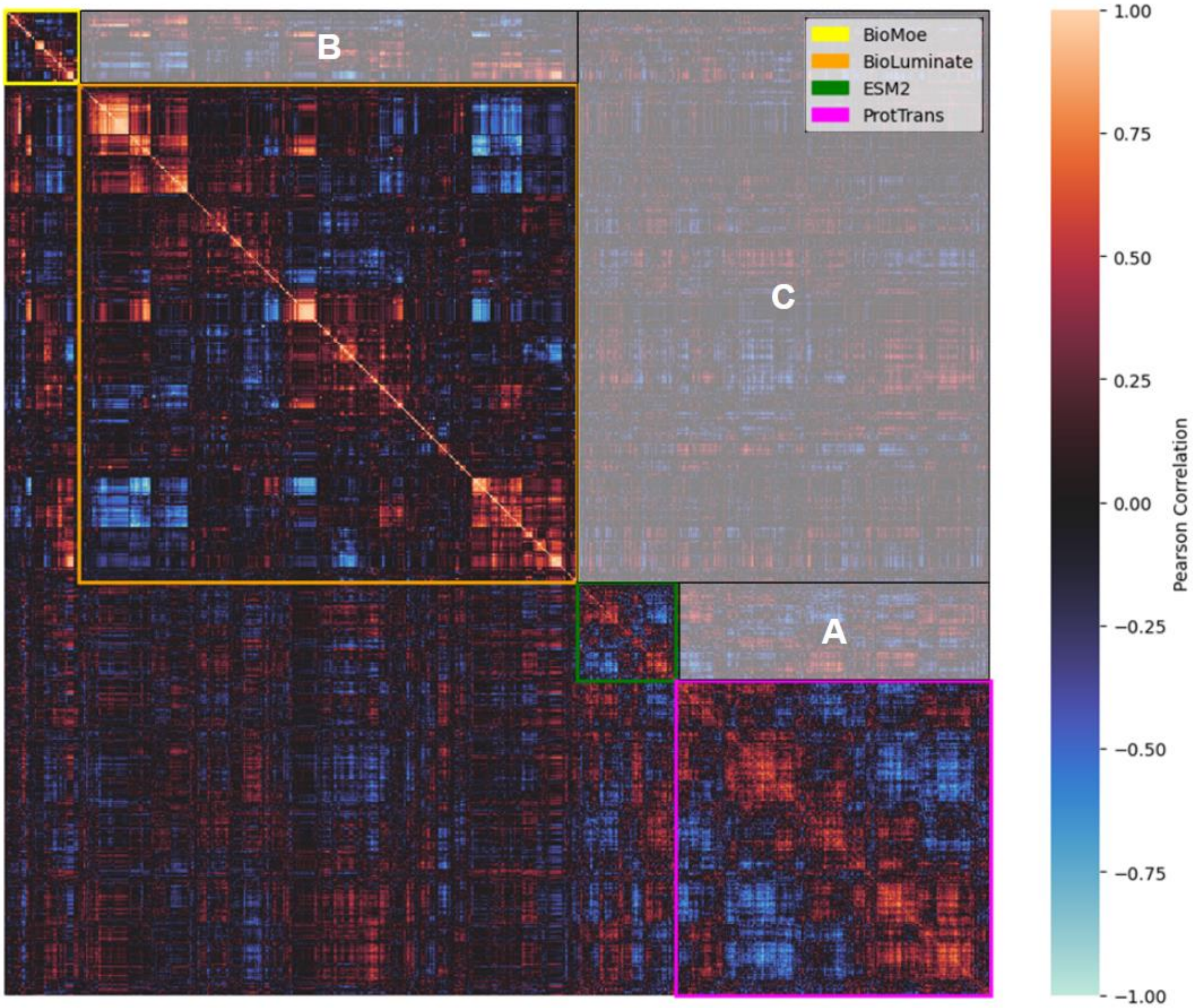


*Figure S7.1 – Feature Correlation Clustermap for All Descriptor Sets.*

The correlation clustermap visualization illustrates covariance between groups of descriptors. Matching row/column indices refer to the same descriptor. The figure is symmetrical across the diagonal (y = -x). Color indicates the Pearson's Correlation between a pair of descriptors. Feature order across the axis is determined by agglomerative clustering features first by descriptor type, and then by Pearson correlation. Colored boxes indicate descriptor sets. The features within each box indicate correlation between descriptors within a given descriptor set. Areas outside of the boxes indicate correlation between different descriptor sets.

In region A, the significant correlation between the two protein language model embeddings (ESM-2, ProtTrans) indicates that these two descriptor sets are learning similar molecular properties. Similarly, in Region B, significant feature correlation is seen between structural descriptor sets (BioMOE, BioLuminate). In region C, off-axis correlation between protein language model embeddings and structure-based descriptors (BioMOE, BioLuminate) suggest that protein language models are learning biophysical molecular properties.

The presence of significant clusters within and across all feature sets indicates significant feature redundancy. These analyses illustrate the challenges of feature multicollinearity in the descriptors sets, and reinforce the necessity of robust feature selection pipelines to reduce the risk of overfitting.

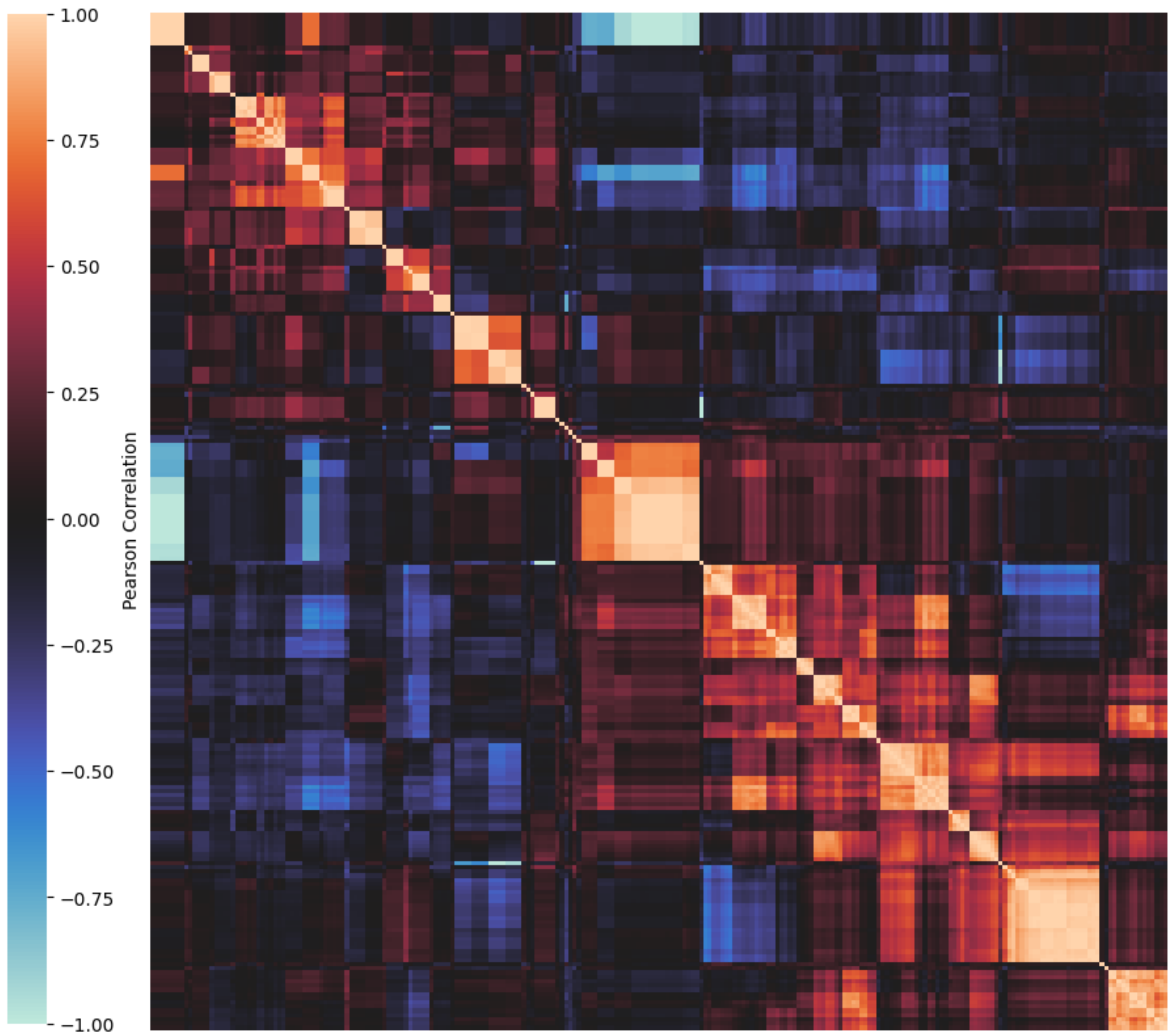


*Figure S7.2 – BioMOE Descriptor Correlation Clustermap*

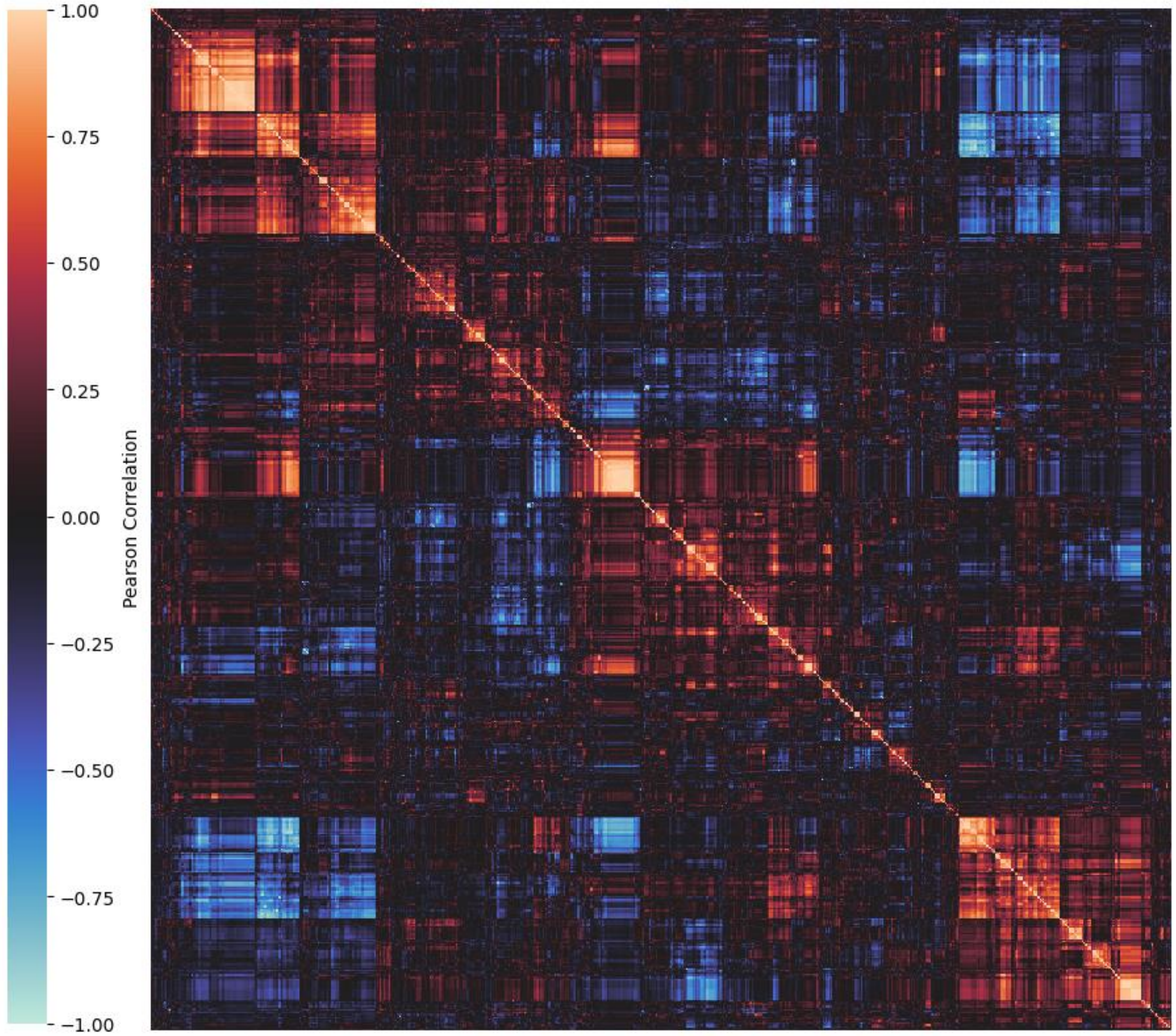


*Figure S7.3 – BioLuminate Descriptor Correlation Clustermap*

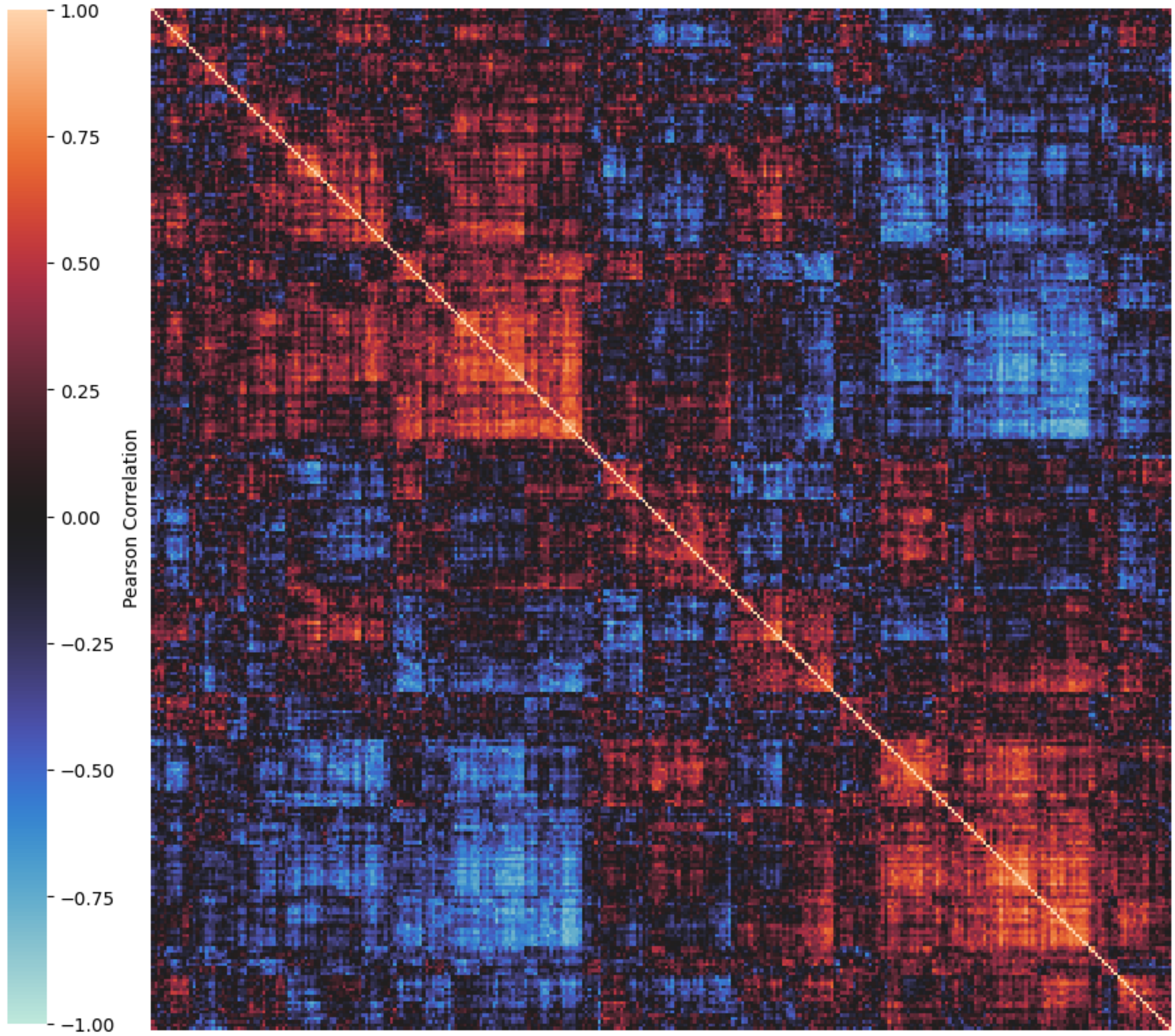


*Figure S7.4 – ESM2 Descriptor Correlation Clustermap*

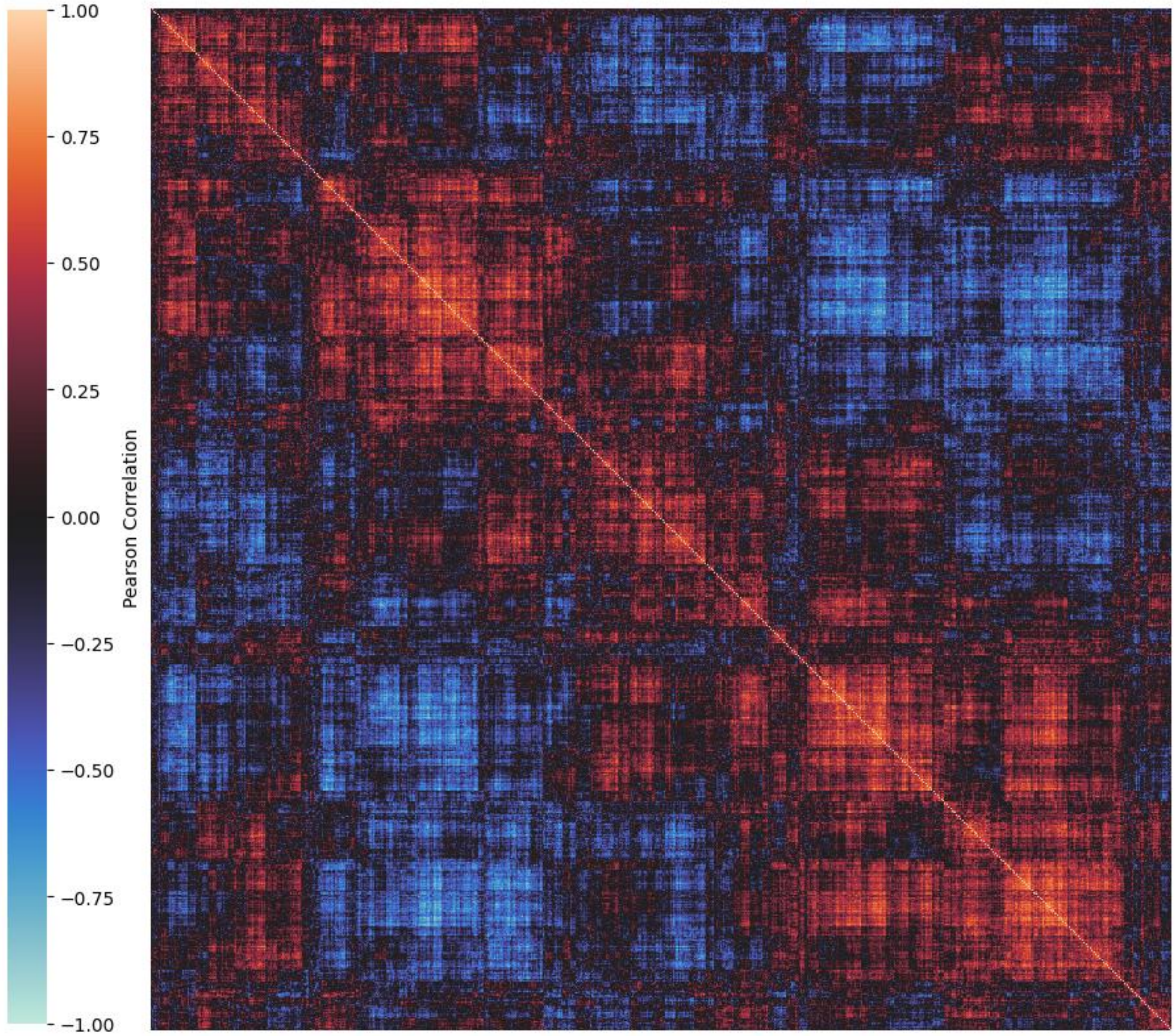


*Figure S7.5 – ProtTrans Descriptor Correlation Clustermap*

#### Section F) Model Feature Importance

Figure S8 illustrates relative feature importance of features in each model. Model feature importance is compared based on SHAP values, a model-agnostic metric for the contribution of features to model predictions derived from game theory. SHAP values assess the marginal importance of features by considering all possible combinations, interactions, and dependencies between features. However, computing SHAP values for large datasets can be computationally intensive. To reduce computational burden, the training set is summarized as 10 clusters using K-means clustering, feature importance is calculated for each cluster, and the mean absolute value feature importance is reported.

For chromatography resins in NaOAc (Figures S8.1 to S8.4), the majority of features with high SHAP values are descriptors of protein charge, which suggests these resins are operating primarily as ion exchangers in this salt system. The top cationic resins (CEX and MMCEX) both include descriptors related to positive charge patches, zeta potential, and formal charge. Meanwhile the anionic resins (AEX and MMAEX) both include descriptors related to negative charge patches. For AEX and CEX in NaOAc the combination of structure-based biophysical and sequence-based protein language model descriptors improved the $Q^2$ by 0.05 and 0.22 respectively over BioLuminate descriptors alone. For CEX in NaOAc, model SHAP values for protein language model-based descriptors are relatively low with only one sequence descriptor appearing in the top ten features. However, for AEX in NaOAc sequence descriptors are of higher importance, accounting for four of the top ten features.

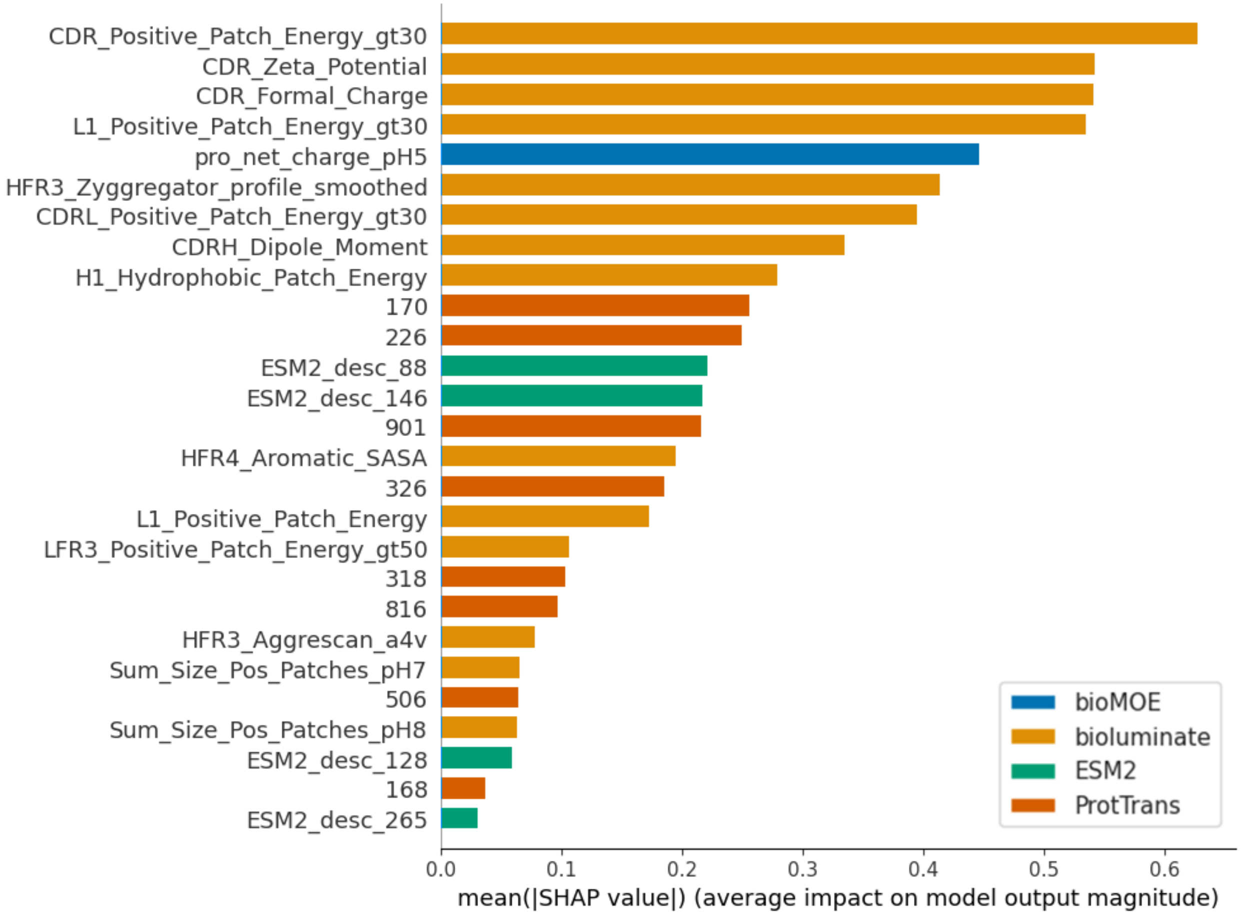


*Figure S8.1 – SHAP feature importance values for CEX in NaOAc*

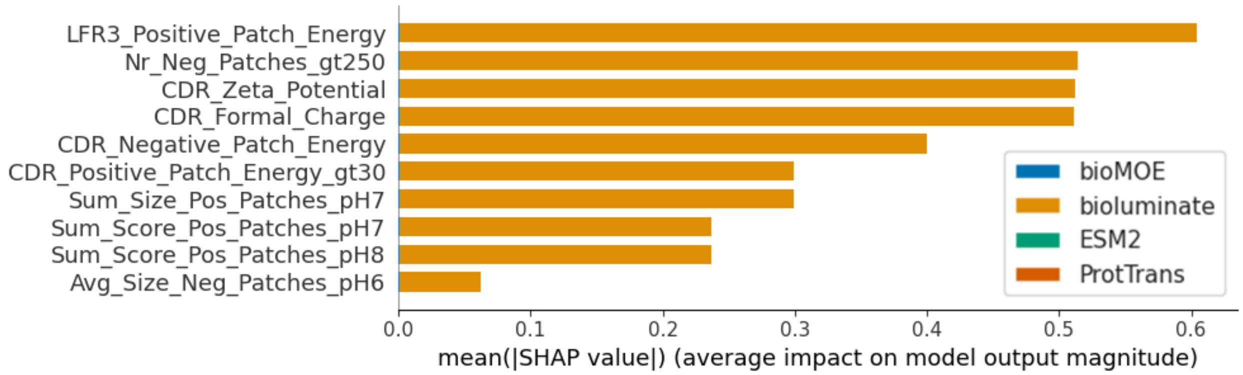


*Figure S8.2 – SHAP feature importance values for MMCEX in NaOAc*

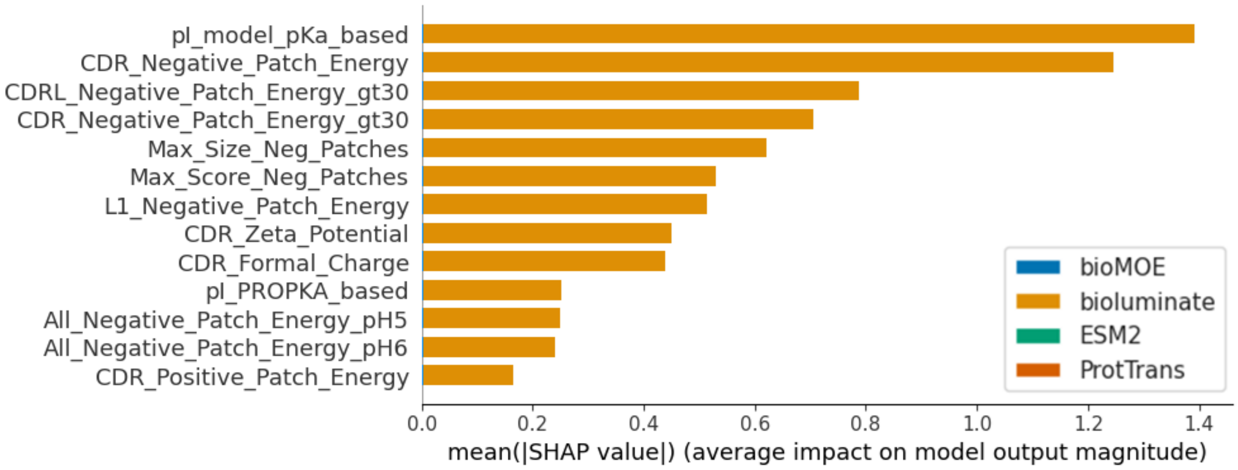


*Figure S8.3 – SHAP feature importance values for MMAEX in NaOAc*

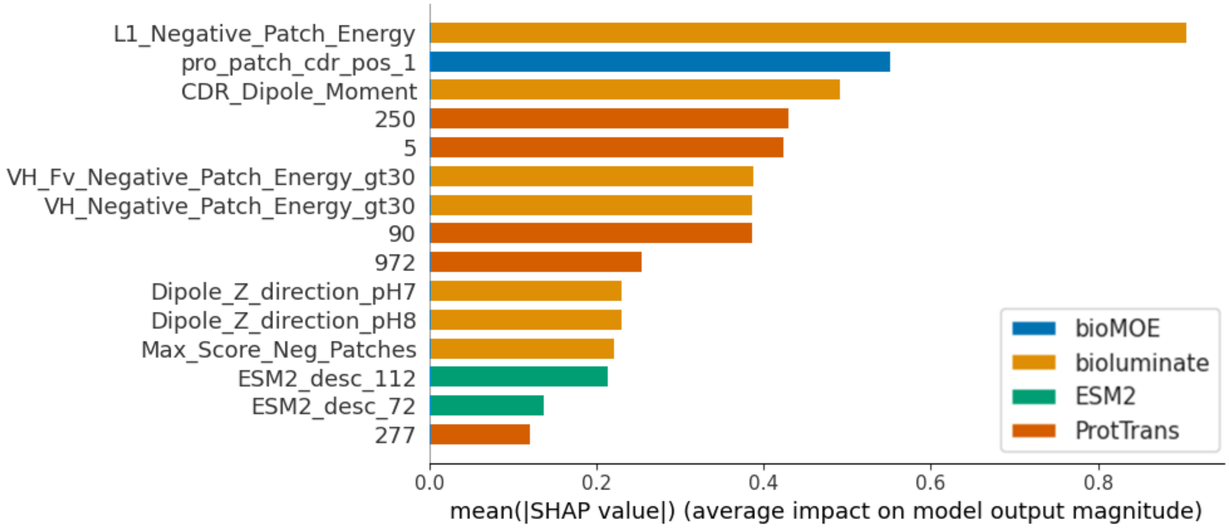


*Figure S8.4 – SHAP feature importance values for AEX in NaOAc*

Similar to MMCEX in NaOAc, MMCEX in $Na_2SO_4$ (Figure S8.5) also primarily relies on descriptors of charge patches and formal charge, but also has high SHAP feature importance values for hydrogen bond acceptor solvent accessible surface area. This observation suggests that in the presence of $Na_2SO_4$ the carboxyl functional group in the MMCEX ligand may be participating in hydrogen bonding, possibly forming cyclic acetic acid dimers with the side chains of aspartic and glutamic acid or bonding to amino acids with nitrogenous side chains.

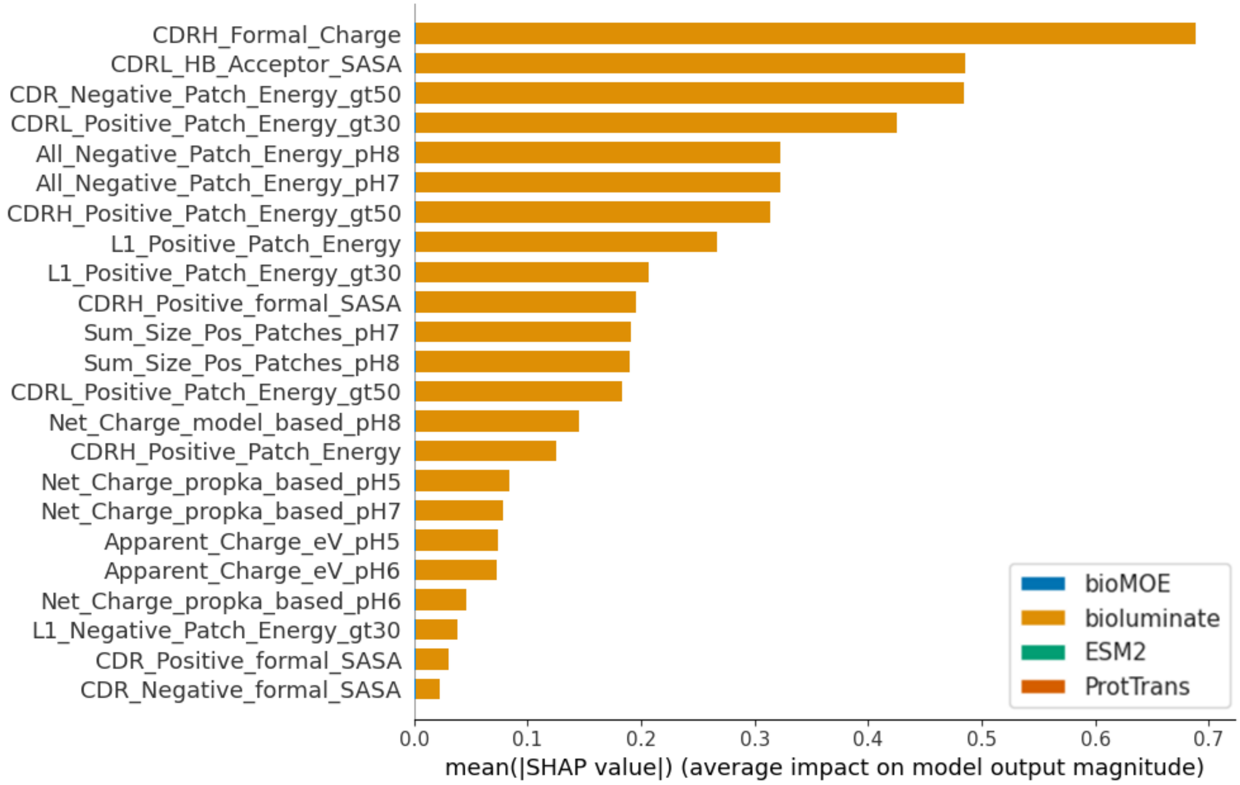


*Figure S8.5 – SHAP feature importance values for MMCEX in $Na_2SO_4$*

The most important features by SHAP value for HIC in $Na_2SO_4$ are shown in Figure S8.6. Notably eight of the top ten features with highest SHAP values for hydrophobic binding including the most important descriptor are sequence-based descriptors from protein language models. While sequence-based protein language model descriptors alone do not produce good predictions ($Q^2$ of ~0.2), the high SHAP values attributed to sequence-based descriptors suggest that they may provide complementary information relevant to prediction of hydrophobic interaction chromatography. Protein language models are thought to learn properties of amino acids and protein sequences, such as hydrophobicity, that are relevant to proper protein folding from co-evolutionary information in large sequence datasets (see citations in section 1.3 of the main text).

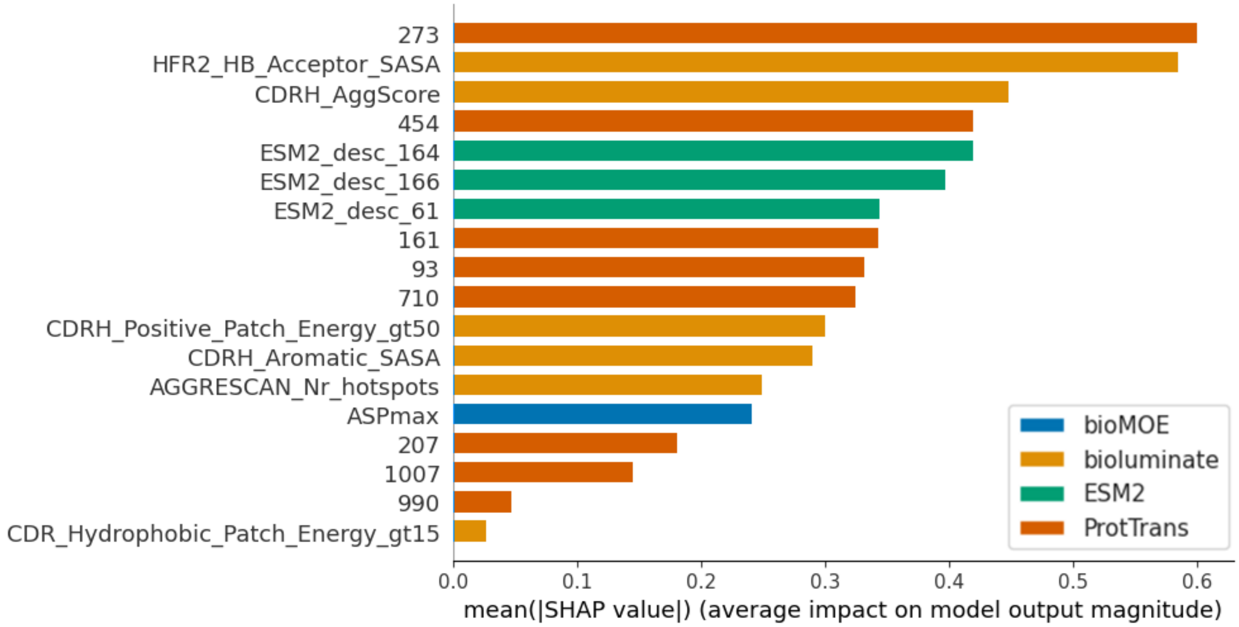


*Figure S8.6 – SHAP feature importance values for HIC in $Na_2SO_4$*

The most important features by SHAP value for MMAEX in $Na_2SO_4$ are shown in Figure S8.7. By far the most important feature is protein ultraviolet (UV) absorbance at 280 nm wavelength (i.e pro_coeff_280). UV absorbance at this wavelength is primarily due to aromatic rings in the sidechains of tryptophan and tyrosine in the protein. This feature was not selected for the MMAEX resin in NaOAc for which binding, as previously discussed, is predicted predominantly by charge-based descriptors. This feature was also not selected for the HIC resin in $Na_2SO_4$, whose ligand has an aromatic ring but lacks the quaternary amine moiety seen in the MMAEX ligand. This observation suggests that there is a hydrophobic interaction between the aromatic rings in the protein and the positively charge quaternary amine on the chromatography ligand. Therefore, the high SHAP values observed for UV280 may suggest that $Na_2SO_4$ may promote electrostatic cation-π interactions between the protein's solvent accessible aromatic amino acid sidechains and the ligands' cationic quaternary amine moieties. There may also be some relatively minor contribution from π- π stacking between the protein's solvent accessible aromatic amino acid sidechains and the ligands' phenyl moieties due to van der Waals forces. This analysis provides insight into the mechanisms of binding for MMAEX in NaOAc vs $Na_2SO_4$, and helps explain potential differences in selectivity of separations in these two salt species.

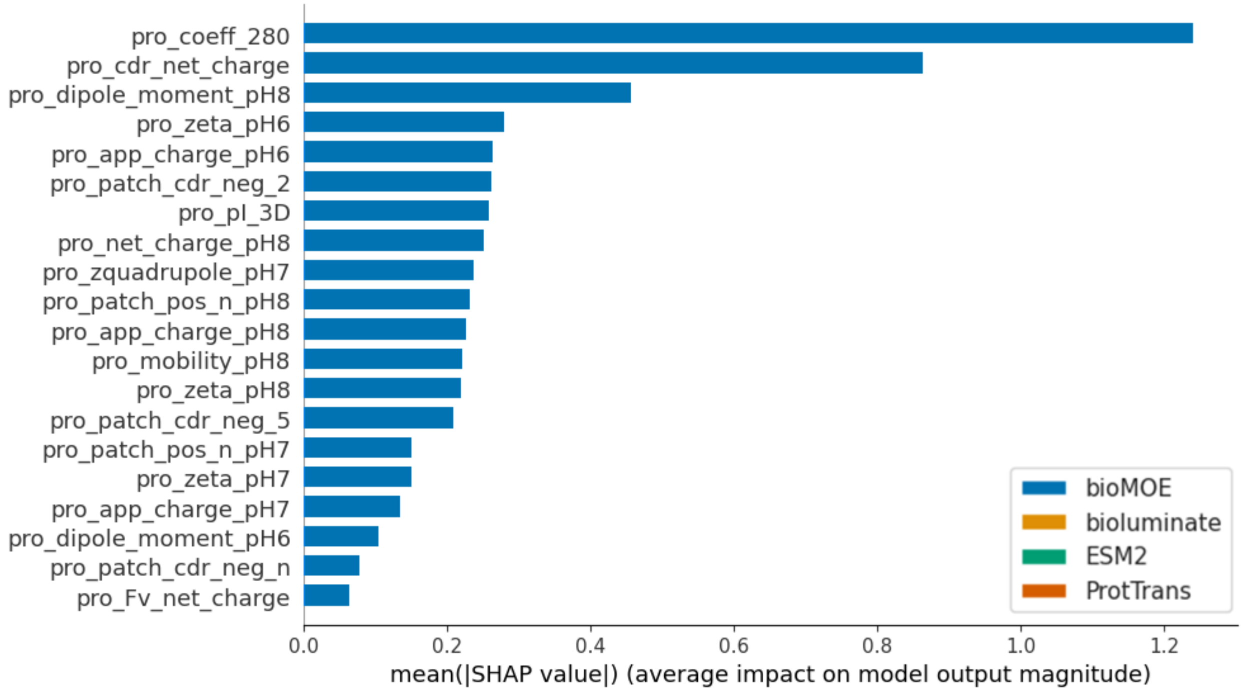


*Figure S8.7 – SHAP feature importance values for MMAEX in $Na_2SO_4$*
